# Analysis of eukaryotic lincRNA sequences reveals signatures of repressed translation in species under strong selection

**DOI:** 10.1101/737890

**Authors:** Anneke Brümmer, Rene Dreos, Ana Claudia Marques, Sven Bergmann

**Affiliations:** Department of Computational Biology (DBC), University of Lausanne, Lausanne, Switzerland; Center for Integrative Genomics (CIG), University of Lausanne, Lausanne, Switzerland; Swiss Institute of Bioinformatics (SIB), Lausanne, Switzerland; Department of Integrative Biomedical Sciences, University of Cape Town, Cape Town, South Africa

**Author notes:** equal last authors (Ana Claudia Marques, Sven Bergmann).

**Keywords:** lincRNA, tRNA abundance, codon usage bias, computational sequence analysis, ribosome binding, tRNA adaptation index, translation regulation, cell type-specificity, cytoplasmic localization, small protein-encoding ORF

## Abstract

Long intergenic non-coding RNAs (lincRNAs) represent a large fraction of transcribed loci in eukaryotic genomes. Although classified as non-coding, most lincRNAs contain open reading frames (ORFs), and it remains unclear why cytoplasmic lincRNAs are not or very inefficiently translated.

Here, we analysed signatures of repressed translation in lincRNA sequences from six eukaryotes. In species under stronger selection, i.e. fission yeast and worm, we detected significantly shorter ORFs than in intronic and non-transcribed control regions, a suboptimal sequence context around start codons for translation initiation, and trinucleotides (“codons”) corresponding to less abundant tRNAs than codons in control regions, potentially impeding translation elongation.

We verified that varying tRNA expression levels affect ribosome-binding to lincRNAs by analyzing data from five human cell lines. Notably, for three cell lines, codons in abundant cytoplasmic lincRNAs corresponded to lower expressed tRNAs than control codons, substantiating cell type-specific repression of lincRNA translation in human. Finally, comparing non-coding with peptide-encoding ORFs suggested that codon usage at the start of ORFs to be of particular relevance for ribosome-binding.

The identified sequence signatures may assist distinguishing peptide- from real non-coding lincRNAs in a cell.

## Introduction

Long intergenic noncoding RNAs (lincRNAs) form a functionally heterogeneous class of RNAs longer than 300 nucleotides and lacking protein coding potential (Frankish et al., 2019; Ulitsky and Bartel, 2013). Despite being classified as non-coding, most lincRNAs contain open reading frames (ORFs) flanked by start and stop codons. Although, many small ORFs encoding functional peptides within annotated human lincRNAs have been identified recently (Chen et al., 2020; Martinez et al., 2020; Ouspenskaia et al., n.d.), for the majority of lincRNAs peptide products have not been detected and the mechanisms counteracting translation are unclear.

Differences between coding and non-coding RNAs have been investigated previously. By analysing sequencing of ribosome protected fragments (Ribo-Seq) data, differences in ribosome interaction patterns have been reported, including in the tri-nucleotide periodicity of binding (Calviello et al., 2016; Ji et al., 2015) or in ribosome release (Guttman et al., 2013). Furthermore, discriminating sequence features between human mRNAs and lincRNAs (Niazi and Valadkhan, 2012) and between lincRNAs with and without ribosome-association have been analysed, in human and mouse (Wang et al., 2017; Zeng and Hamada, 2018). These studies identified a poor start codon context of lincRNA ORFs for translation initiation and reported cell type-specific associations between human lincRNAs and ribosomes. Yet, these studies considered only a relatively small set of lincRNAs in only one or two species, and provided little insight into how translation of lincRNAs is hindered.

mRNA translation to proteins is known to be regulated at initiation and during elongation (Eraslan et al., 2019; Nieuwkoop et al., 2020; Riba et al., 2019; Tuller et al., 2010a). While the RNA sequence and secondary structure around start codons are important for translation initiation, it has been demonstrated that the codon usage is a regulator of translation elongation. Specifically, the rate at which a codon is translated correlates with the abundance of its decoding tRNA (Dana and Tuller, 2014). Consequently, mRNAs composed of codons corresponding to more abundant cellular tRNAs tend to be translated more efficiently. Evidence for this mechanism to increase translation efficiency has been observed in several contexts, e.g. under cellular stress conditions, during proliferation and meiosis, and in cancer (Goodarzi et al., 2016; Guimaraes et al., 2020; Sabi and Tuller, 2019; Torrent et al., 2018). The strength of the mRNA codon usage bias varies between species, and it is usually more pronounced in species under stronger selection, with a larger population size or a shorter generation time (Subramanian, 2008). Among eukaryotes, the mRNA codon usage bias is strongest in yeast and weaker in species such as human and mouse. In most species, the codon usage bias is stronger for highly expressed mRNAs, likely because the selection pressure is higher on these mRNAs.

Given these well-studied connections between mRNA sequence features and translation efficiency, here we investigate sequence signatures indicative of repressed translation of lincRNAs in six eukaryotes. In species under strong selection, i.e. fission yeast and worm, we found evidence for several such sequence signatures. We further examined the functionality of a biased lincRNA codon usage in reducing ribosome-binding to lincRNAs using experimental data from five human cell lines.

## Results

### Open reading frames in lincRNAs are frequent

To investigate signatures in lincRNAs sequences counteracting translation efficiency, we focussed on six species (human, mouse, fish, fly, worm and fission yeast). These species were selected because they contain a sizable number (>1000) of annotated lincRNA genes in their genomes and cover a range of natural selection pressure, as observed in their mRNA codon usage bias strengths (Subramanian, 2008).

In order to exclude potential biases in the sequences of lincRNAs, we removed lincRNAs that overlapped other genes, repetitive sequences (Jurka et al., 2005), and likely novel coding regions (predicted based on PhyloCSF score (Lin et al., 2011) or identified from Ribo-Seq data, in case of human (Martinez et al., 2020)) (see Methods and Table S1). For each lincRNA, we extracted all ORFs longer than 30 nucleotides (=10 codons), starting with a canonical start codon (AUG) and ending at the first in-frame stop codon (UAG, UAA, UGA). As a control, we analysed ORFs in randomly selected intronic and intergenic regions of the same length and with the same G+C content (fraction of G and C nucleotides) as lincRNAs (see Methods). These nuclear (intronic) and non-transcribed (intergenic) sequences are not in contact with the translation machinery and therefore provide a reference, against which we evaluated the sequence bias of lincRNAs in each species. We would like to point out that differences in the genomic sequence composition across species may affect the analysed lincRNA measures making them not directly comparable between species. Therefore, we only compare the deviation of lincRNA from control measures (within each species) across species.

We found that most lincRNAs (>93% for each species) had at least one ORF (Figure 1A). The fraction of lincRNAs with ORFs in fission yeast was significantly smaller than that in intronic regions, but ORFs were more often found in lincRNAs than control regions for other species. While the number of ORFs per lincRNA was similar to that in control sequences for fission yeast, significantly more ORFs were found in lincRNAs than in control regions for other species. The median number of ORFs per lincRNA in different species ranged between 5 (fly and worm) and 9 (fission yeast) (Figure S1A).

**Figure 1:**
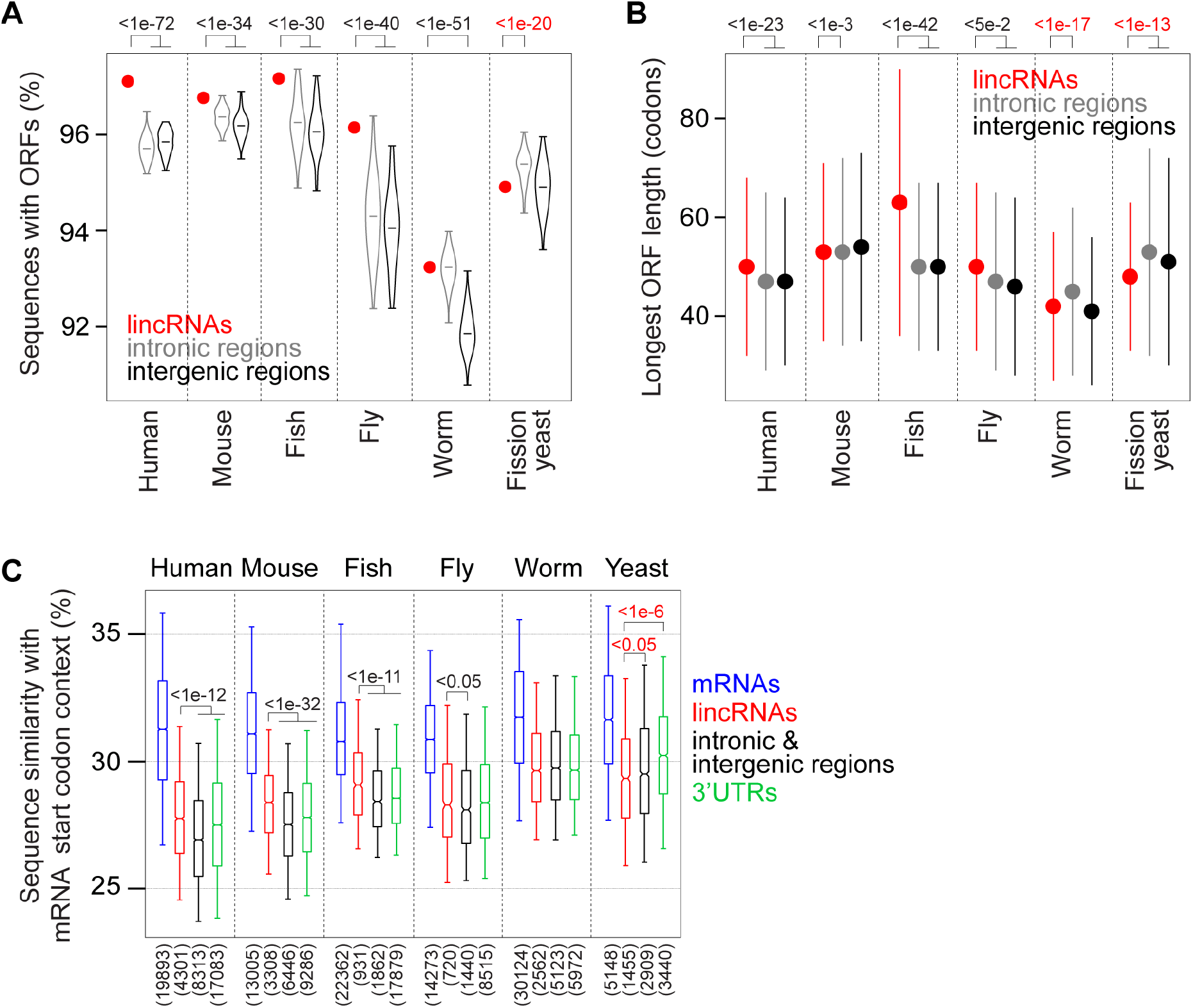
Prevalence, length, and start codon context of lincRNA ORFs. (A) Percentage of lincRNA transcripts with ORFs (>10 codons; red dot) and percentage of random control regions with ORFs (in introns shown as gray and in intergenic regions shown as black violins), for six species. For each lincRNA transcript 10 length- and G+C content-matched control regions were randomly selected in introns and intergenic regions. P values are indicated from a one-sample t test. (B) Median length of longest ORFs in lincRNAs (red), and intronic (gray) and intergenic (black) control regions. Error bars represent median absolute deviations. P values are indicated from Wilcoxon’s rank-sum test. (C) Sequence similarity with the consensus mRNA sequence motif for the region +/−12 nucleotides around start codons (see Methods) for mRNA coding regions (blue), lincRNA longest ORFs (red), intronic and intergenic control ORFs (black), and longest ORFs in 3’UTRs (green). P values (<0.05) are indicated from Wilcoxon’s rank-sum test to compare lincRNAs with control regions or 3’UTRs. P values are marked red, if the median lincRNA value is below the value for control regions.

In the following, we consider for each lincRNA its longest ORF (median length between 44 to 65 codons in different species; Figure 1B) and its first ORF (median length between 21 to 27 codons; Figure S1B). We reasoned that these ORFs might exhibit the strongest signatures of repressed translation, because synthesis of a longer peptide would consume more resources (in case of longest ORFs) and first ORFs would be most exposed to scanning ribosomes starting from the 5’ end of transcripts. Both, longest and first ORFs, were significantly shorter in lincRNAs than these respective ORFs in intronic and intergenic control regions for fission yeast, and than in intronic regions for worm (Figures 1B and S1B). In contrast, in vertebrates, longest and first ORFs in lincRNAs were longer than these respective ORFs in control regions. In all species, mRNA coding regions were much longer than longest lincRNA ORFs (5.6 to 9.1 times for median lengths).

In summary, lincRNA ORFs are rarer and shorter compared to ORFs in control regions in fission yeast and to some extent in worm, but not in other species.

In the following, for each mRNA we only consider the isoform with the longest coding region and for each lincRNA gene the isoform with the longest ORF (Table S1), or the isoform with the longest among first ORFs. As control ORFs, we selected from random intronic and intergenic regions of the same length as the lincRNA isoform harboring the longest ORF, the longest ORF (>10 codons) with G+C content matching that of the longest lincRNA ORF. Additionally, we analysed longest ORFs in 3’ untranslated regions (UTRs) of coding genes, which likely represent the least translated cytoplasmic RNA sequences (Guttman et al., 2013), and first ORFs in 5’ UTRs of coding genes, which were reported to resemble ribosome association with that of lincRNAs (Chew et al., 2013).

### RNA sequence around lincRNA start codons appears suboptimal for translation initiation in yeast and worm

Since the RNA sequence and structure around start codons was identified as a regulator of translation in mRNA (Eraslan et al., 2019), we examined if the start codon context of lincRNA ORFs would be less optimal for initiating translation. We found that in all species the sequence context around lincRNA start codons was less specific than the consensus mRNA sequence motif, which showed similarity with the Kozak sequence motif (Kozak, 1989) (Figure S2A). For fission yeast, the sequence around lincRNA start codons was less similar to the consensus mRNA sequence motif than that of control ORFs and longest ORFs in 3’UTRs (Figure 1C). For worm, there was no difference in similarity between lincRNAs and controls, and for vertebrates and fly, we found a higher similarity with the consensus mRNA sequence motif for lincRNAs than for controls. In the case of first ORFs in lincRNAs, we found less similarity with the consensus mRNA sequence motif than for first ORFs in control regions for worm, and than for first ORFs in 5’UTRs for human and mouse (Figure S2B).

To examine this further, we asked whether the similarity with the consensus mRNA sequence motif would be reduced for conserved (evolutionary older) lincRNAs compared to younger lincRNAs, since older lincRNAs may have had more time to adapt their sequences. We grouped lincRNAs according to their level of conservation for species, where suitable conservation data were available (not for yeast and fish; see Methods). Indeed, for worm, the similarity with the consensus mRNA sequence motif was lower for more conserved lincRNAs than for less conserved lincRNAs, potentially indicating that older transcripts developed stronger signatures for inefficient translation initiation (Figure S2C). There was no significant difference between lincRNAs with different conservation levels for fly, mouse and human.

In summary, for yeast lincRNAs and conserved lincRNAs in worm the RNA sequence around lincRNA start codons appears to be less favourable for translation initiation, but not for other species.

We next analysed the RNA structure around start codons in different species (Figure S3). We found that in human and mouse this region had a higher minimum free energy, calculated using RNAfold (Hofacker, 2004) (see Methods), and thus was more accessible in lincRNAs and control regions than in mRNAs. This could be a consequence of a higher G+C content in mRNAs in these species (Haerty and Ponting, 2015; Niazi and Valadkhan, 2012). In contrast, in fly and fission yeast, the structural accessibility was reduced in lincRNAs compared to mRNAs, which may hinder an interaction between ribosomes and lincRNA start codons in these species. Overall, it appears that the regulation by the RNA secondary structure around translation start sites is complex and might differ between species.

### Codon composition of lincRNA ORFs is distinct from mRNA coding regions, and from control ORFs in worm and yeast

The codon usage of mRNA is biased and it contributes to translation regulation (Hanson and Coller, 2018; Tuller et al., 2010b). We investigated if biased frequencies of trinucleotides, which for simplicity we refer to as codons, in lincRNA ORFs can contribute to counteracting translation efficiency. To gain initial insight into the characteristics of codon usage in different RNA types, we performed a multiple correspondence analysis of their codon counts in different species (Table S2). We found a clear separation between RNA types in the first two components space (Figures 2A and S4A). While for vertebrates and fly, lincRNA codon composition was positioned in between those of mRNAs and control ORFs, but much closer to the latter, lincRNA and mRNA codon usages were located on opposite sides relative to controls for worm and fission yeast, in the first two components space. 3’UTR codon usage tended to be more distant from the codon usage of mRNAs than the codon usage of lincRNAs (especially for fish, worm and yeast), while the codon usage of first ORFs in 5’UTRs was closest to mRNAs for human, mouse and worm. These distinctions in codon usage between RNA types were reflected in the patterns of correlation strengths between codon frequencies of different RNA types (Figure 2B). In particular, for worm and yeast, the correlation between mRNA and control codon frequencies was stronger than the correlation between lincRNA and mRNA codon frequencies. First and longest ORFs of lincRNAs had very similar codon frequencies (r^2^>0.89 in all species; Figure S2A). Therefore, we decided to focus on analysing the codon usage of longest ORFs in lincRNAs from hereon.

**Figure 2:**
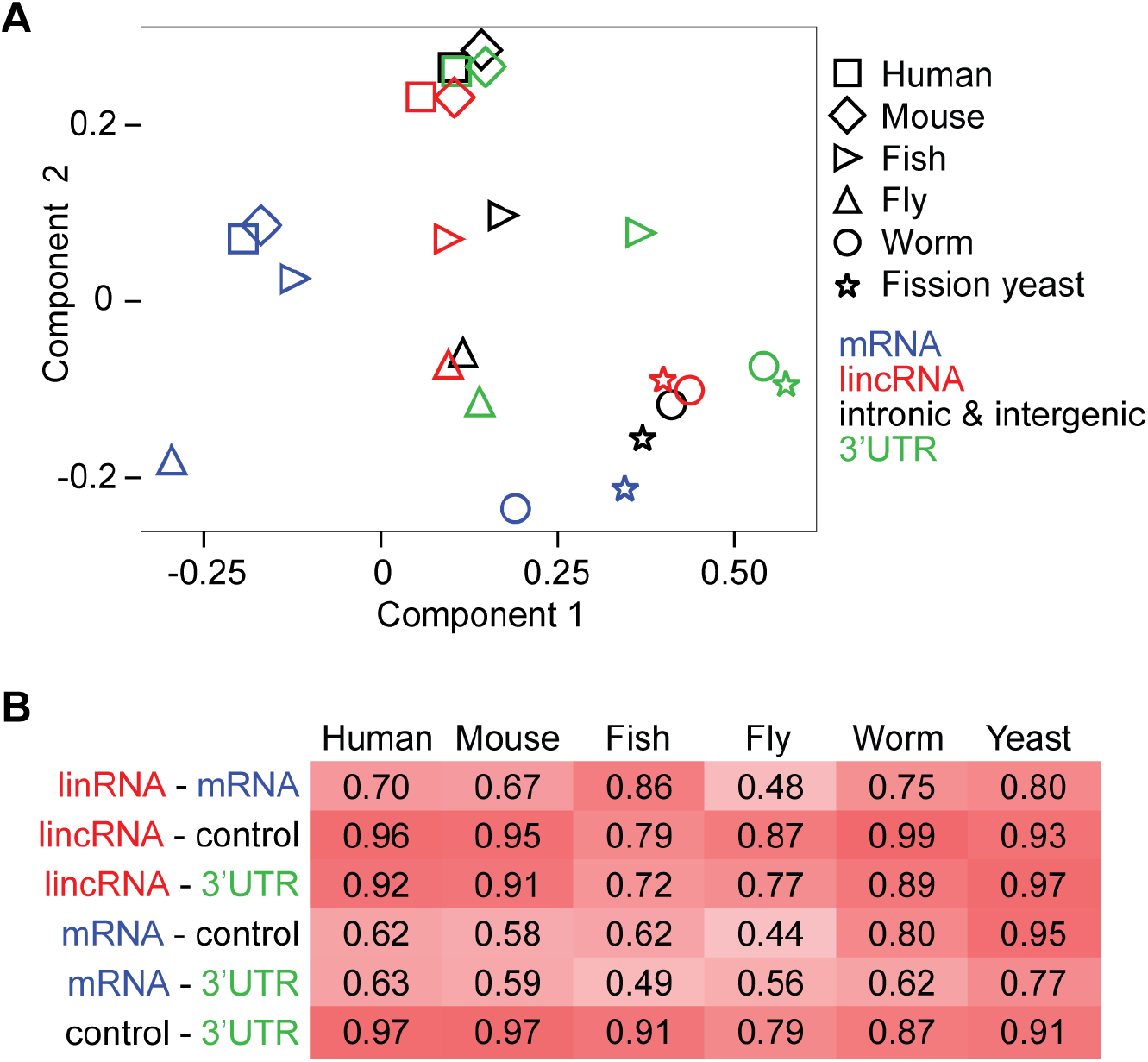
Comparison of trinucleotides (“codons”) in mRNA and longest ORFs in lincRNAs, control regions and 3’UTRs. (A) First two components space from a multiple correspondence analysis performed on trinucleotide (“codon”) counts (excluding start and stop codons) in mRNA coding regions (blue) and longest ORFs in lincRNAs (red), intronic and intergenic control regions (black), and 3’UTRs (green) for six species. (B) Spearman correlation coefficients between codon frequencies for the same ORFs as in (A) and for six species. Corresponding p values are given in Table S4.

To rule out that differences in G+C content between RNA types (Haerty and Ponting, 2015) are responsible for the observed differences in codon usage, we separately analysed codon counts among groups of codons with identical G+C content (Figure S4B). For most G+C content codon groups, we found a separation between codon usage of mRNAs, lincRNAs and controls in the first two components space of a multiple correspondence analysis. Importantly, for yeast and worm, lincRNA and mRNA codon usages were located on opposite sides from controls in the first two components space for all G+C content codon groups. We conclude that differences in G+C content do not drive the observed differences in codon composition between RNA types.

In summary, lincRNA codon usage is distinct from mRNA codon usage and closer to that of control regions, in most species. In species with strong mRNA codon usage bias (worm and fission yeast), lincRNA and mRNA codon usages appear to have diverged in opposite directions relative to the codon composition of control regions.

### lincRNA codons correspond to less abundant tRNAs than mRNA codons, and than codons in control regions for species under strong selection

Next, we investigated how codon usages of different RNA types relate to tRNA abundances. As a first estimate of the tRNA abundances in different species we used the number of annotated tRNA genes for each tRNA anticodon type (see Methods), which has been shown to correlate well with tRNA abundances (Tuller et al., 2010a). We used previously determined wobble-base pairing and tRNA editing efficiencies (dos Reis et al., 2004) to estimate effective tRNA anticodon abundances for all codons including those that lack a complementary tRNA encoded in the genome (see Methods). We found that mRNA codon frequencies correlated better with relative tRNA abundances than codon frequencies of lincRNAs, for all species (Figure 3A). For yeast and fly, this discrepancy was more pronounced among highly expressed cytoplasmic mRNAs and lincRNAs (see Methods). For the two species with the strongest mRNA codon usage bias, worm and fission yeast, lincRNA codon frequencies correlated less with tRNA abundances than control codon frequencies, suggesting a lincRNA codon usage bias towards codons corresponding to lower abundance tRNAs in these species. In fly, the correlations between tRNA abundances and codon frequencies was similar for lincRNAs and controls, and lincRNAs were better correlated with tRNA abundances than controls in human, mouse and fish.

**Figure 3:**
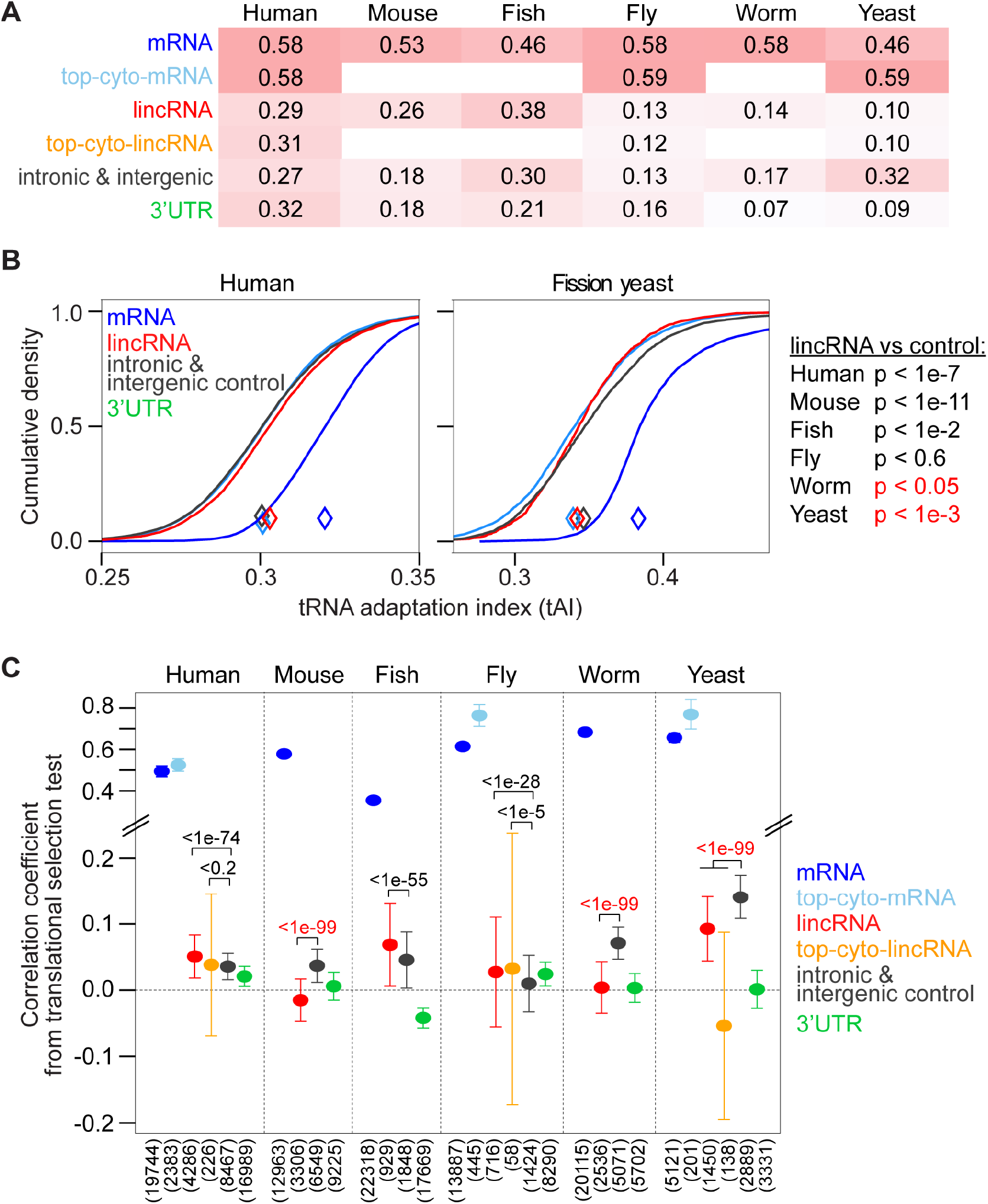
Correspondence between tRNA abundances and codon frequencies in different RNA types. (A) Spearman correlation coefficients between effective tRNA anticodon frequencies based on tRNA gene copy numbers (see Methods) and codon frequencies in mRNA coding regions and longest ORFs in lincRNAs, intronic and intergenic control regions, and 3’UTRs for six species; and for codon frequencies in cytoplasmic mRNAs and lincRNAs for human, fly and yeast (see Methods). Corresponding p values are given in Table S4. (B) Cumulative density of tRNA adaptation indexes (tAIs) for mRNA coding regions (blue), and longest ORFs in lincRNAs (red), intronic and intergenic control regions (black) and 3’UTRs (green) for human (left panel) and fission yeast (right panel). Similar plots for other species are shown in Figure S5A. On the right, p values from Wilcoxon’s rank-sum test to compare tAIs of lincRNAs with those of control regions are indicated for all six species. P values are marked red, if the median tAI of lincRNAs is lower than that of control regions. (C) Translational selection test (adapted from dos Reis et al. (dos Reis et al., 2004); see Methods): Pearson correlation coefficient between tAI and the adjusted effective number of codons, for ORF types and species as in (A). Error bars indicate 90% confidence intervals estimated from 1000 times bootstrapping. P values are indicated from Wilcoxon’s rank-sum test for the comparison of bootstrapped values from lincRNAs with those of controls. P values are marked red, if the correlation for lincRNAs is lower than that for control regions.

Previously, the tRNA adaptation index (tAI) was defined as a measure for the correspondence between codon frequencies in an ORF and tRNA abundances (dos Reis et al., 2003). The tAI ranges from 0 to 1, where higher values indicate a preferential usage of codons decoded by more abundant tRNAs. We calculated the tAI to examine the overall correspondence between codon and tRNA frequencies (as opposed to the correspondence of frequencies among synonymous codons; see Methods). We found that, for all species studied, mRNA coding regions had significantly (p<1e-300, Wilcoxon rank-sum test) higher tAI values than longest ORFs in lincRNAs (Figures 3B and S5A). This was unrelated to the difference in number and length of mRNA coding regions and lincRNA ORFs, since for subsampled mRNAs, accounting for number and lengths of lincRNA ORFs, we also found significantly larger tAIs than for lincRNAs (data not shown). Compared to ORFs in control regions, lincRNA ORFs had significantly lower tAIs in worm and fission yeast, while for vertebrates, tAIs were higher for lincRNA ORFs than for control ORFs (Figure 3B and Figure S5A). These tAI differences between lincRNAs and control ORFs reflect the results from the correlation analysis (Figure 3A), potentially indicating that lincRNA ORFs have adapted to use codons corresponding to less abundant tRNAs in yeast and worm. Notably, longest ORFs in 3’UTRs also had lower tAIs than control ORFs in yeast, worm and fly, but similar tAIs than control ORFs in human, mouse and fish.

To further evaluate the extent and direction of codon usage bias in lincRNAs, we related tAIs to those of randomized ORFs. In particular, we compared with tAIs for trinucleotides in shuffled ORFs, preserving the nucleotide content, and in frame-shifted ORFs, preserving the nucleotide content and sequence, including potential functional RNA sequence or structure motifs (see Methods; Figure S6). To account for underlying (di-)nucleotide biases in the genomic sequence of each species, we performed the same randomizations with ORFs in control regions and used this comparison as a reference. Overall, tAI differences (ΔtAI) between original and shuffled ORFs or between original and frame-shifted ORFs were positive for mRNAs in all species, indicating generally higher tAIs for the original coding sequences. ΔtAIs between original and shuffled ORFs were mostly also positive for other ORF types, including control regions, which might be due to dinucleotide biases in the genomic sequences. ΔtAIs between original and frame-shifted ORFs were closer to zero for other ORF types. For worm and yeast, ΔtAIs were significantly lower for lincRNAs relative to ΔtAIs controls, for comparison with shuffled and frame-shifted ORFs. This indicates that tAIs of lincRNA ORFs tended to be lower than those of their randomized ORFs more often than this was the case for control ORFs. In vertebrates, ΔtAIs for lincRNAs were overall larger than those of controls for the comparison with shuffled ORFs, and for comparison with frame-shifted ORFs in mouse. Together, the comparison with tAIs of randomized ORF sequences supports that lincRNA ORFs exhibit a codon composition that corresponds to less abundant tRNAs compared to control ORFs, in worm and yeast. In particular, the comparison with frame-shifted ORFs, which alter the codon composition but maintain the nucleotide sequence, strengthens the hypothesis that lincRNA ORFs are biased for preferential usage of codons corresponding to less abundant tRNAs as opposed to for maintaining specific RNA sequence or structure motifs, in these species.

A test for translational selection of mRNA codon usage in a species, was proposed by dos Reis et al. (dos Reis et al., 2004). In particular, the correlation between tAI and a measure for the deviation of the codon usage from an equal usage of synonymous codons, while accounting for the G+C-bias at the third codon position, was proposed. Here, we adapted this to test for a correlation between tAI and the overall codon usage bias (not just among synonymous codons; see Methods). Using this correlation test, we confirmed that correlation coefficients were larger for mRNAs than for lincRNAs (r>0.35 and r<0.10, respectively, for all species), indicating a stronger adaptation of mRNA codon usage to tRNA abundances than for lincRNA codons (Figure 3C). Furthermore, the correlation was larger among cytoplasmic mRNAs (r>0.52) in human, fly and yeast, while it tended to be smaller among cytoplasmic lincRNAs (r<0.04) in human and yeast. Strikingly, for mouse, worm and yeast, the correlation coefficients for lincRNAs (and cytoplasmic lincRNAs in yeast) were significantly lower than those for control ORFs (p<1e-99), and correlations were also significantly lower for 3’UTR ORFs compared to control ORFs for all species (p<1e-52, Wilcoxon’s ranksum test with correlation coefficients from bootstrapped samples; see Methods). This indicates that codons in non-coding ORFs are less adapted to tRNA abundances than control codons in these species, potentially to counteract efficient translation.

### Codons in cytoplasmic lincRNAs correspond to lower expressed tRNAs than control codons in three out of five human cell lines

In multicellular eukaryotes, tRNA expression is often tissue- and cell type-specific (Dittmar et al., 2006; Pinkard et al., 2020). We utilized this to investigate the impact of varying tRNA abundances on ribosome-binding to cytoplasmic lincRNAs. We focussed on five human cell lines (GM12878, HEK293, HeLa-S3, HepG2, and K562), for which extensive experimental data are available to quantify relative tRNA expression levels, total and cytoplasmic lincRNA expression levels, and ribosome-binding to lincRNAs ((Aktaş et al., 2017; Calviello et al., 2016; Cenik et al., 2015; Consortium and The ENCODE Project Consortium, 2004; Huang et al., 2019; Kishore et al., 2013; Martinez et al., 2020; Solomon et al., 2017; Subtelny et al., 2014); see Methods and Table S3).

As tRNA gene counts are not cell type-specific, various types of experimental data were used previously to estimate cell type-specific tRNA abundances, such as H3K27ac-ChIP-Seq (Gingold et al., 2014) and smallRNA-Seq data (Hernandez-Alias et al., 2020; Shi et al., 2018). For human HEK293 cells, data is also available from two dedicated experimental methods to quantify tRNA abundances, DM-tRNA-Seq (Zheng et al., 2015) and hydro-tRNA-Seq (Gogakos et al., 2017). We evaluated the concordance of different methods in HEK293 and found that smallRNA-Seq-based tRNA quantifications agree better with those from dedicated experimental approaches than H3K27ac-ChIP-Seq-based tRNA quantifications (Figure S7A; see Methods). Therefore, we used smallRNA-Seq data to quantify cell type-specific tRNA abundances in human cell lines.

We found varying tRNA abundances between cell lines (Figure 4A), with GM12878, HeLa-S3 and K562 having relatively similar tRNA abundances (Spearman correlation >0.87), while those in HEK293 and HepG2 were more different from other cell lines (Spearman correlation <0.75). Using these tRNA abundances we calculated cell line-specific tAIs (Figure 4B), and confirmed that abundant cytoplasmic lincRNAs had significantly lower tAIs than cytoplasmic mRNAs in all cell lines. tAIs of cytoplasmic lincRNA were also lower than tAIs of mRNAs with cytoplasmic expression levels matching those of lincRNAs (p<1e-12; see Methods). For three out of five cell lines (GM12878, HeLa-S3 and K562), tAIs of abundant cytoplasmic lincRNAs were significantly lower than tAIs of control ORFs (p<0.03) and than tAIs of all expressed lincRNAs in a cell line (p<0.04). For HEK293 and HepG2, tAIs of highly expressed cytoplasmic lincRNAs were not different from those of all expressed lincRNAs or control ORFs. For GM12878, HeLa-S3 and HEK293, a small negative correlation between tAI and cytoplasmic expression level of lincRNAs is detectable (Figure S8A).

**Figure 4:**
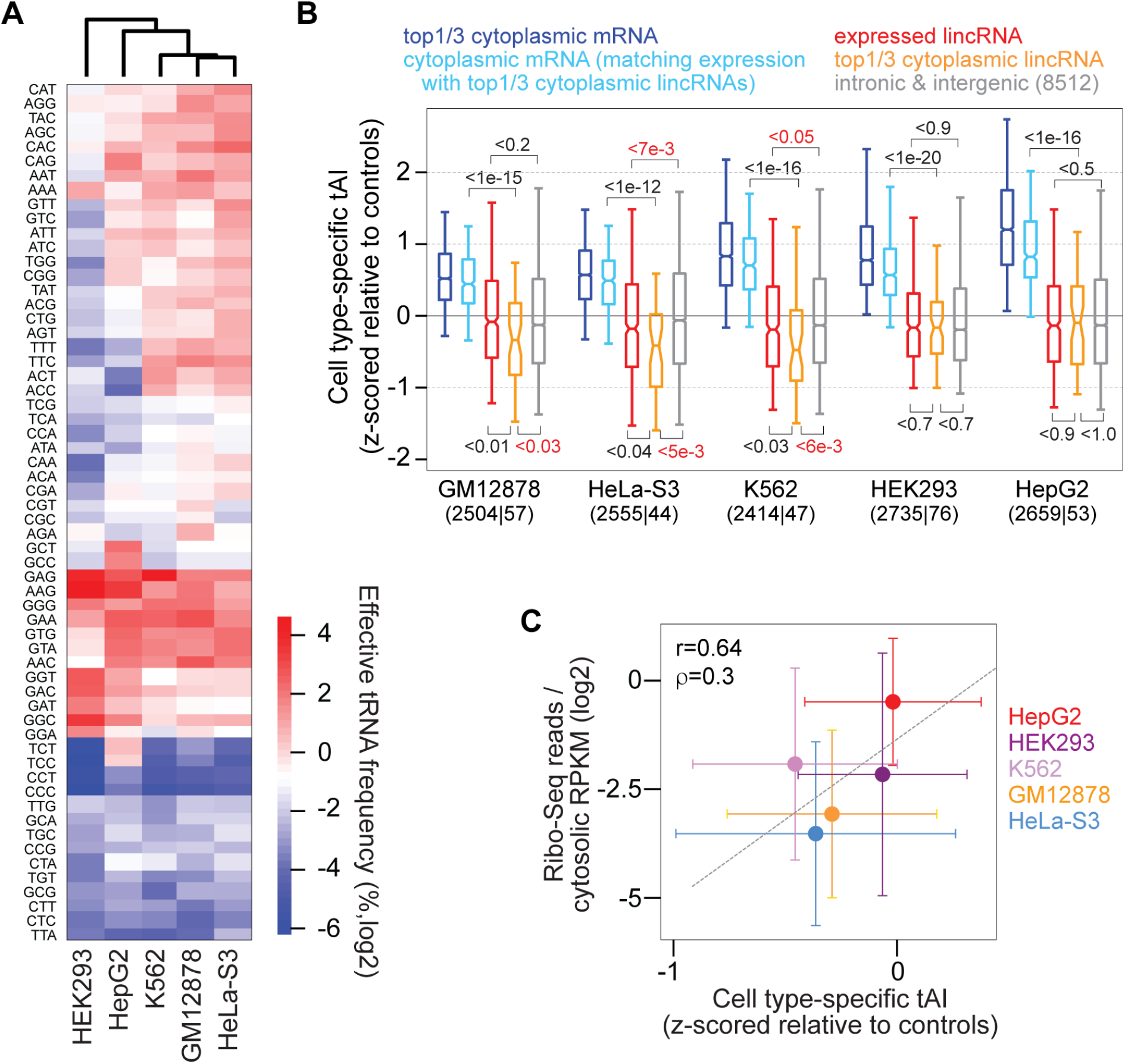
Cell type-specific tAIs and ribosome-binding in five human cell lines. (A) Clustering (based on Euclidean distance) of effective tRNA anticodon frequencies, estimated from smallRNA-Seq data, in five human cell lines (indicated at bottom). (B) Boxplots of cell type-specific tAIs (z-scored relative to those of intronic and intergenic control ORFs) for top-1/3 cytoplasmic expressed mRNAs (dark blue), mRNAs with cytoplasmic expression levels matching those of top-1/3 cytoplasmic lincRNAs (light blue), expressed lincRNAs (red), top-1/3 cytoplasmic expressed lincRNAs (orange) and controls (gray), for five human cell lines (label at bottom). The numbers of top-1/3 cytoplasmic mRNAs and lincRNAs are indicated in parentheses at bottom. P values are calculated using Wilcoxson’s rank sum test. (C) Relationship between cell type-specific tAIs (z-scored relative to control tAIs) and relative ribosome-binding (see Methods) for 41 lincRNAs that are cytoplasmic in all five human cell lines. Dots represent median values and error bars median absolute deviations. Pearson (r) and Spearman (ρ) correlation coefficients between median values are indicated.

We next focussed on lincRNAs that we classified as cytoplasmic in all five human cell lines (see Methods). We confirmed that tAIs for these 41 lincRNAs vary between cell lines, with significantly higher values in HepG2 and HEK293 than in K562 (p<0.07, Wilcoxon’s rank sum test). These differences in cell line-specific tAIs were largely reflected in ribosome-binding differences (estimated from Ribo-seq data; see Methods) between cell lines for the same 41 cytoplasmic lincRNAs (Figure 4C). A relationship between cell type-specific tAI and ribosome-binding was also detectable when comparing lincRNAs that are cytoplasmic in any pair of cell lines (Figure S8B). In particular, ribosome-binding in HepG2 was significantly higher than in other cell lines (p<0.04, Wilcoxon’s rank sum test). Notably, increased translation of lincRNAs in liver and kidney was reported before (van Heesch et al., 2019).

In summary, the analysis of human cell lines revealed tAI differences for cytoplasmic lincRNAs between cell lines. tAIs of abundant cytoplasmic lincRNAs were smaller than those of control ORFs in three out of five cell lines, suggesting reduced ribosome-binding.

### Ribosome-binding reflects tAI differences between lincRNAs and small protein-encoding RNAs, particularly for codons at ORF starts

To understand the relationship between tAI and ribosome binding further, we focussed on three types of cytoplasmic RNAs: mRNAs, lincRNAs, and annotated lincRNAs with small protein-encoding ORFs (smORFs; see Methods). We observed that tAIs of lincRNAs and smORFs are significantly different from those of mRNAs in K562 (Figure 5A) and other human cell lines (Figure S9A). This difference in tAIs was concordant with differences in ribosome binding, estimated from Ribo-seq data, in two ways. First, the proportion of transcripts with Ribo-seq reads was significantly larger for mRNAs than for smORFs, and both of these were larger than the fraction of lincRNA ORFs with Ribo-seq reads (Figures 5B and S9B). Second, relative ribosome-binding (calculated as the ratio of normalized Ribo-seq read counts to cytoplasmic expression level of the ORF-harboring transcript; see Methods), which accounts for differences in ORF lengths and expression between RNA types, was significantly lower for lincRNAs than for other cytoplasmic RNA types, in K562 (Figure 5C) and most human cell lines (Figure S9C).

**Figure 5:**
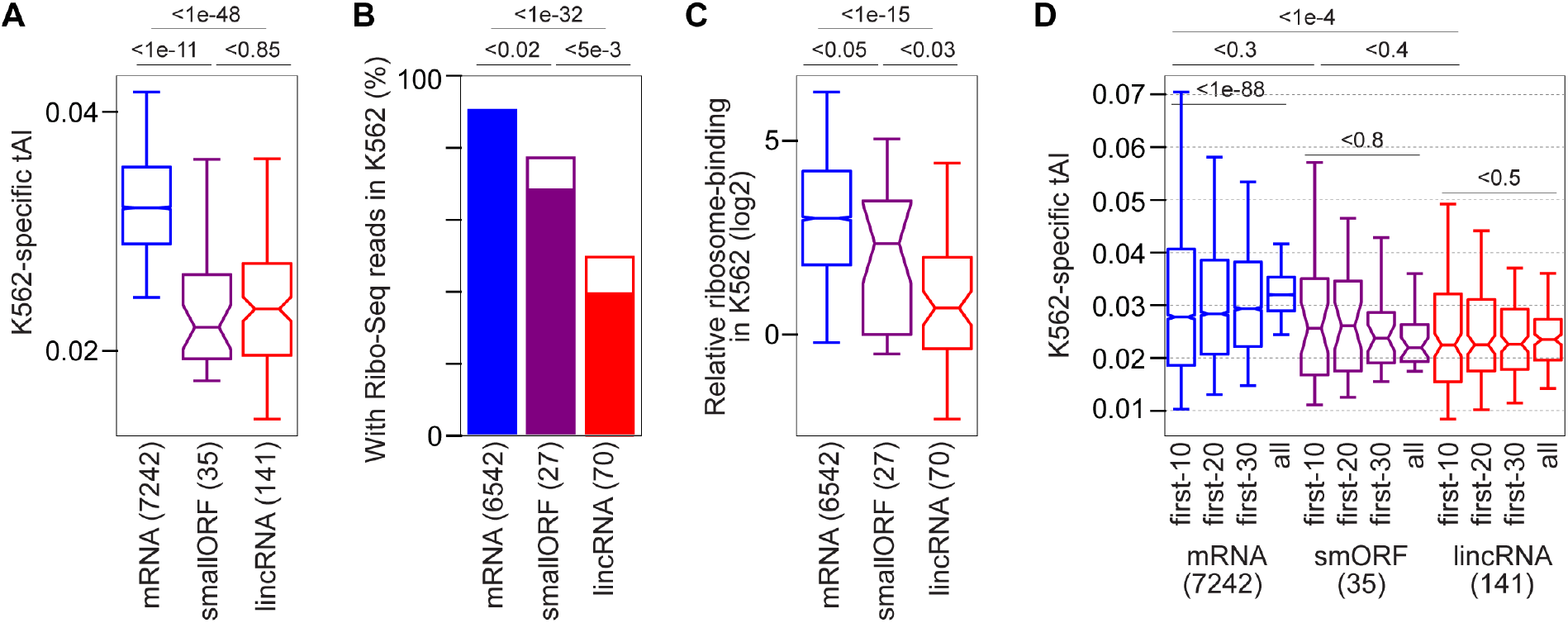
tAIs and ribosome-binding to coding and lincRNA ORFs in K562 cells. (A) Cell type-specific tAIs of three types of cytoplasmic RNA in K562 cells: mRNAs (blue), annotated lincRNAs with small protein-encoding ORFs (smORFs; purple) and lincRNAs (red). P values are indicated from Wilcoxon’s rank-sum test. The number of RNAs of each type is indicated in parentheses at bottom. (B) Fraction of ORFs from (A) with in-frame Ribo-Seq reads (filled bars: with more than one read, empty bars: with one read). P values are indicated from Fisher’s exact test to compare fractions of RNAs with at least one in-frame Ribo-Seq read (number of RNAs indicated in parentheses at bottom). (C) Relative ribosome-binding (calculated as the ratio of normalized in-frame Ribo-Seq reads to the cytoplasmic expression level; see Methods) for ORFs with Ribo-Seq reads. P values are indicated from Wilcoxon’s rank-sum test. Number of RNAs of each type is indicated in parentheses at the bottom. (D) Cell type-specific tAIs at ORF beginnings: the first 10, 20 or 30 codons downstream of start codons (first three boxes for each RNA type), and entire ORFs (last box for each RNA type). P values are indicated from Wilcoxon’s rank-sum test. The number of RNAs of each type is indicated in parentheses at bottom.

Despite this concordance, the differences in relative ribosome-binding between lincRNAs and smORFs were more pronounced than their differences in tAIs, for most cell lines (Figures S9A and C). Previously, the codon usage immediately downstream of mRNA start codons was found to be distinct and was proposed to facilitate efficient translation initiation and elongation (Bentele et al., 2013; Tuller et al., 2010a). Thus, we investigated tAIs calculated for the first codons downstream of start codons in different RNA types. Indeed, tAIs calculated for the first 10 or 20 codons (first-10-codon or first-20-codon tAIs) tended to be higher for smORFs compared to tAIs of entire smORFs, and first-10-codon or first-20-codon tAIs of smORFs were more similar to those of mRNAs (Figures 5D and S9D). There was no such trend for lincRNA tAIs. Notably, we found no indication for the RNA sequence or structure context around start codons of smORFs and lincRNA ORFs to explain the observed ribosome-binding differences between these cytoplasmic RNA types (Figures S9E and F).

These observations indicate that cytoplasmic lincRNAs have lower cell type-specific tAIs than cytoplasmic, small-protein coding RNAs, and support an association with reduced ribosome-binding to cytoplasmic lincRNAs.

## Discussion

In the absence of any functional role of the peptides resulting from lincRNA translation, it would likely be advantageous to reduce stable associations between lincRNAs and ribosomes for several reasons: unwanted lincRNA translation wastes energy (Wagner, 2005), reduces the pool of ribosomes available for mRNA translation (Raveh et al., 2016), and may lead to the synthesis of peptides with possibly harmful interference. Furthermore, it may hinder a potential regulatory function of lincRNAs, and, given the association between mRNA translation and transcript stability (Presnyak et al., 2015; Tuck et al., 2020), translation of lincRNAs might impact their cytoplasmic expression levels, with potentially disadvantageous consequences.

Here, we analysed lincRNA sequences from six eukaryotes to seek for signatures of repressed or inefficient translation (summarized in Figure 6). The analyzed signatures comprised general properties, such as the abundance and lengths of ORFs in lincRNAs, signatures related to translation initiation, such the sequence and secondary structure around start codons, and signatures related to efficient translation elongation, such as the codon usage and its relation with tRNA abundances. While we found evidence for almost all analysed signatures for lincRNAs in fission yeast and worm, in the other species the various signatures were similar or less present in lincRNAs compared to intronic and intergenic control regions.

**Figure 6:**
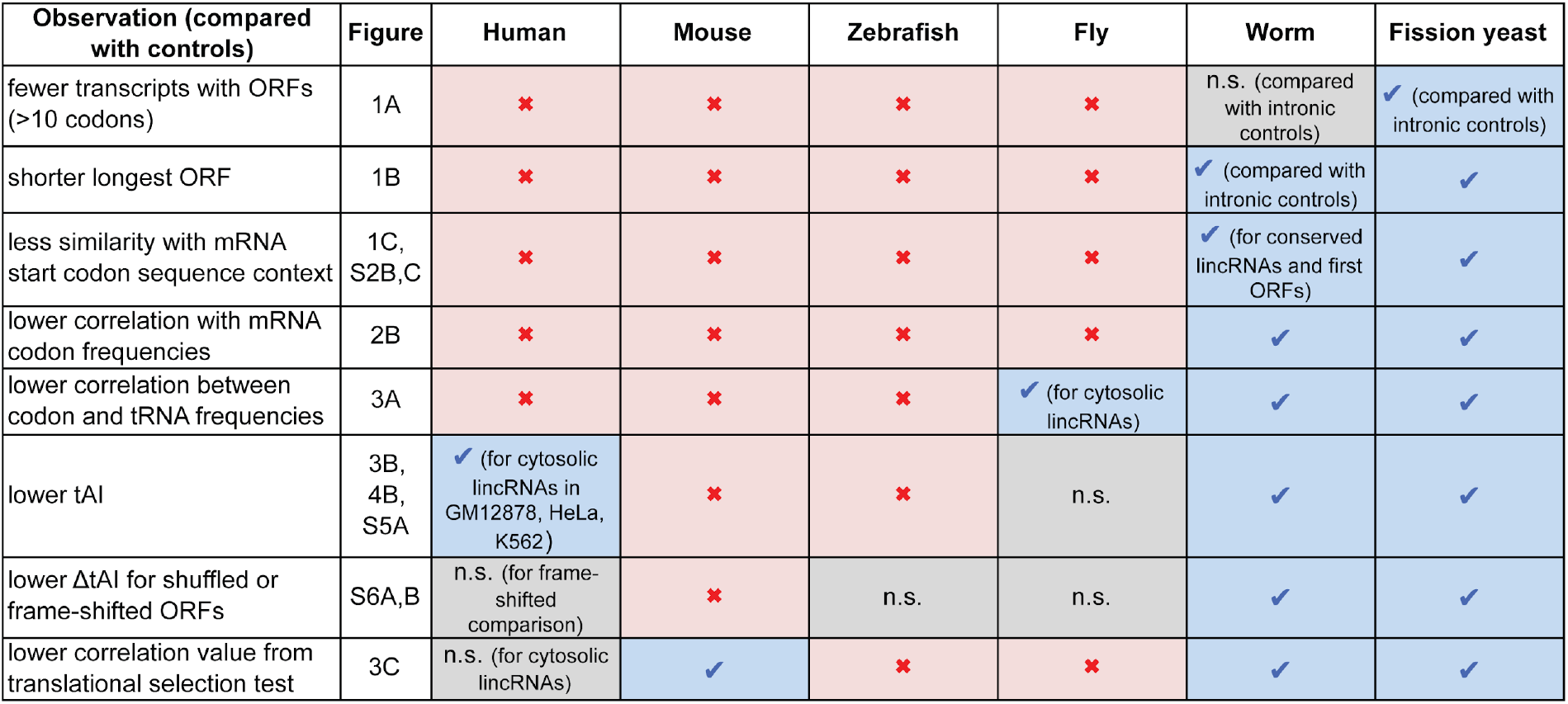
Summary of the observed signatures of repressed lincRNA translation in six species. Blue check marks indicate that the corresponding measure is significantly lower for lincRNAs compared to intronic and intergenic control regions, n.s. stands for not significant, and a red cross indicates that the corresponding measure is significantly higher for lincRNAs compared to controls.

A reason for these differences across species might be that selection pressure is stronger in fission yeast and worm than in other species. Thus, adaptation of lincRNA sequences to counteract translation would likely also be stronger in yeast and worm than in other eukaryotes, where also translational selection on mRNA was found to be weaker (dos Reis and Wernisch, 2009).

We acknowledge that the observed effect sizes of the signatures counteracting translation in fission yeast and worm appear small in some cases. However, the consistency of the effects across multiple, largely independent signatures substantiates the hypothesis that lincRNA sequences in fission yeast and worm are biased to counteract translation. Besides that, due to lower expression levels likely connected with reduced selection pressure on lincRNAs compared to mRNAs, lincRNAs would be expected to have sequence biases with smaller effect sizes than those found in mRNAs.

The lack of observed signatures counteracting translation in species, such as human and mouse, may also suggest that lincRNA translation is not universally repressed, but more in a cell-type specific manner, in these species. Indeed, estimating cell type-specific tRNA expression levels, we observed for three out of five human cell lines that codons in abundant cytoplasmic lincRNAs correspond to less expressed tRNAs than trinucleotides in control ORFs (Figure 4B). This is consistent with previous reports of cell type- or condition-specific translation of human lincRNAs (Chen et al., 2020; Martinez et al., 2020; Ouspenskaia et al., n.d.; van Heesch et al., 2019; Wang et al., 2017), potentially because certain peptides cause no harm or actually serve a function in certain cell types or under certain conditions.

Interestingly, we detected similar signatures counteracting translation in longest ORFs of 3’UTRs, and the translational selection test revealed significantly lower correlation values for 3’UTRs than for controls in most species (except fly; Figure 3C). Several observations were even slightly stronger for 3’UTRs than for lincRNAs, such as the fact that codon usage was more different from that in of mRNAs, for worm, yeast and fish (Figure 2A), or tAIs were lower, for worm, yeast and fly (Figure S5A). This could be the result of a stronger selection on 3’UTRs than on lincRNA ORFs, potentially because 3’UTRs are mostly higher expressed in the cytoplasm and are evolutionarily older (Ulitsky and Bartel, 2013), and thus had more time for adapting their sequence to tRNA abundances. On the other hand, the fact that tAIs of 3’UTRs for vertebrates were not lower than control tAIs (Figure S5A), may indicate that the selection pressure on non-coding ORFs in these species is not strong enough to result in tAIs lower than for control ORFs, or that cell-type specific translation regulation is more relevant in these species.

Notably, the codon usage of first ORFs in 5’UTRs was more similar to that of mRNAs than to lincRNAs, for human, mouse and worm, likely indicating a frequent functional role of upstream ORFs in these species.

Functionally, we validated an impact of varying tRNA expression levels on ribosome-binding to lincRNAs by analyzing cell type-specific tAIs and ribosome-binding to cytoplasmic lincRNAs in five human cell lines. We propose a mechanistic link between the tAI of a lincRNA ORF and its ribosome-binding, that would allow a cell type-specific repression of lincRNA translation. Interestingly, tAIs for the first 10-20 codons of lincRNA ORFs and small protein-encoding ORFs appeared to better match differences in their ribosome-binding, in most cell lines, suggesting a particular importance of codons at the beginnings of ORFs for ribosome engagement and translation (or its avoidance in case translation would be disadvantageous).

In conclusion, in this study we provided a comprehensive analysis of signatures counteracting efficient translational in lincRNA sequences of six species and in five human cell lines. The identified sequence signatures may help in distinguishing bona-fide lincRNAs with regulatory roles in the cytoplasm from those coding for peptides. An interesting aspect is how cell type- and condition-specific expression of tRNAs and cytoplasmic lincRNAs imposes different constraints on the evolution of lincRNA sequences to either curb ribosome-binding in certain cell types, or promote it to enable peptide translation in others. Analysis of the impact of natural genetic variation or targeted mutations in lincRNA sequences on ribosome-binding and peptide translation might further shed light onto these questions.

## Material and Methods

### Gene annotations

Gene annotations and genomic sequences were downloaded from GENCODE (Frankish et al., 2019) (www.gencodegenes.org) for human (v19 corresponding to hg19) and mouse (vM16 corresponding to mm10), from Ensembl (www.ensembl.org) for zebrafish (dr11), fruit fly (dm6) and worm (ce11), and from EnsemblFungi (http://fungi.ensembl.org) for fission yeast (ASM294v2). For worm (*C. elegans*), we additionally included lincRNA genes that were recently identified by Akay et al. (Akay et al., 2019), and were not contained the Ensembl gene annotations. We chose fission yeast as opposed to the more commonly studied budding yeast, because the number of annotated lincRNA genes was much larger in fission yeast (>1000) than in budding yeast (<100). For all species, we excluded genes on mitochondrial chromosomes from our analysis, as these are translated by mitochondrial tRNAs.

### mRNA coding regions

For each mRNA gene we only analyzed the longest coding region starting with a canonical start codon (AUG), from all coding regions in different transcript isoforms.

### Identification of open reading frames (ORFs) in lincRNAs

LincRNA genes were excluded that overlapped (by at least one nucleotide on either strand) with other gene annotations, with regions of high PhyloCSF score (Lin et al., 2011) (PhyloCSF Novel tracks downloaded from https://data.broadinstitute.org/compbio1/PhyloCSFtracks/ for human, mouse, fly, and worm), and, in case of human, with regions that were experimentally identified to encode small proteins based on Ribo-Seq data in human cell lines (Supplementary Table 1 of (Martinez et al., 2020)). Furthermore, lincRNA genes were excluded that overlapped (by more than 30 nucleotides) with simple repeats or low complexity regions (repeat masker annotations downloaded for human, mouse, fish, fly, and worm from http://hgdownload.soe.ucsc.edu/goldenPath/), as such regions may bias the sequence composition of lincRNAs. The number of remaining lincRNAs after each of these filtering steps is listed in Table S1. ORFs longer than 30 nucleotides (=10 codons) that start with a canonical start codon (AUG) and end at the first in-frame stop codon (UAG, UAA, UGA) were identified using a custom python script. For each lincRNA gene the transcript isoform harboring the longest ORF or longest first ORF was considered for further analysis.

### Intronic and intergenic control sequences

As control, we considered nuclear (intronic) and non-transcribed (intergenic) sequences. Intronic regions were taken from mRNA genes and were required to not overlap any exons on either strand and excluding 10 nucleotides flanking exons. Intergenic regions were required to not overlap any gene annotation on either strand. We excluded intronic and intergenic regions that overlapped with likely novel coding regions (based on PhyloCSF novel track for human, mouse, fly and worm, and Ribo-Seq data for human) and with repetitive sequence regions (from repeat masker annotations for human, mouse, fish, fly and worm). For each lincRNA transcript harboring the longest ORF (or the longest first ORF), a region of the same length was randomly selected from intronic and intergenic regions using bedtools shuffle (Quinlan and Hall, 2010). In each random control region, the longest or first ORF (>10 codons) was identified. If the fraction of G and C nucleotides (rounded to two decimal places) equaled the one of the lincRNA ORF, the region was retained as a control region for that lincRNA, otherwise the random selection was repeated until a region with equal G+C content was found.

For the comparison of ORF identification in lincRNAs and control regions (Figures 1A,B and S1A,B), we randomly selected, for each lincRNA transcript, 10 length- and G+C content-matched control regions in intronic and in intergenic regions.

### Small protein-encoding ORFs (smORFs)

In Figure 5, human smORFs were longest ORFs in annotated lincRNAs that overlapped (by at least 1 nucleotide, on the same strand) with a small protein-encoding region, identified experimentally by Martinez et al. in three human cell lines based on Ribo-Seq data (Supplementary Table 1 of (Martinez et al., 2020)), or with a region with high PhyloCSF score (Lin et al., 2011) (downloaded from https://data.broadinstitute.org/compbio1/PhyloCSFtracks/ for human).

### Sequence and structure context around start codons

To analyse the sequence context around start codons the fraction of nucleotides (A,C,G,T) at each position in the region +/− 12 nucleotides around start codons was counted. The information content (*I*) at each position was calculated as: 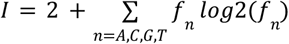, where *f_n_* is the frequency of nucleotide *n*. The probability (*P*) for the consensus mRNA start codon motif (also referred to as similarity) was calculated as: 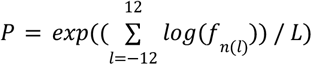, where *L* is the total length of the start codon region and *f_n(l)_* is the frequency of nucleotide *n* (for mRNA) at position *l* around the start codon of an ORF.

For the RNA structure context analysis in Figures S2 and S7E, the minimum free energy of the sequence region +/− 25 nucleotides around start codons was calculated using RNAfold (Hofacker, 2004).

### Conservation scores for lincRNAs and control regions

BigWig files with PhastCons scores were downloaded from the UCSC genome browser (http://hgdownload.soe.ucsc.edu/goldenPath/) for human for multiple alignments of 10 primate genomes to hg19 (primates.phastCons46way.bw), for mouse for multiple alignments of 39 placental vertebrate genomes to mm10 (mm10.60way.phastCons60wayPlacental.bw), for fly for multiple alignments of 26 insect genomes to dm6 (dm6.27way.phastCons.bw), and for worm for multiple alignments of 25 nematode genomes to ce11 (ce11.phastCons26way.bw). The conservation scores for lincRNA transcript isoforms harboring the longest ORFs were calculated as the average PhastCons score over all nucleotides in exons. The conservation scores for intronic or intergenic control regions were calculated as the average PhastCons score over all nucleotides in that region.

### Multiple correspondence analysis

The multiple correspondence analysis of codon counts in ORFs of different RNA types in different species (Figures 2A and S4) was implemented in python.

### Estimation of relative tRNA abundances based on tRNA gene counts

tRNA gene predictions were downloaded from GtRNAdb (http://gtrnadb.ucsc.edu/GtRNAdb2/) (Chan and Lowe, 2016) for all species studied. The number of annotated tRNA genes coding for the same tRNA anticodon type were counted considering high confidence tRNA gene predictions. Effective tRNA abundances were calculated using previously determined weights to account for tRNA editing and wobble-base pairing efficiencies (dos Reis et al., 2004). In particular, the weights *w* are *w*(G:U)=0.41, *w*(I:C)=0.28, *w*(I:A)=0.9999, and *w*(U:G)=0.68, where the first letter denotes the first nucleotide of a tRNA anticodon nucleotide triplet and the second letter the third nucleotide of a codon. Effective tRNA abundances were then calculated as:

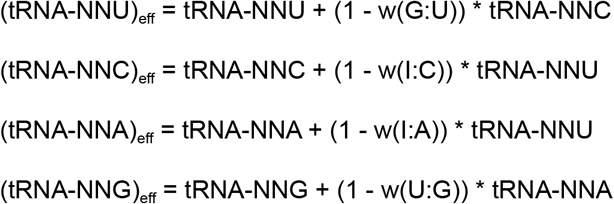

The above nucleotide triplets are the corresponding codon sequences (i.e. the reverse complements of the tRNA anticodon sequences). N stands for any nucleotide.

### Estimation of cell type-specific tRNA abundances

Due to the repetitive nature of tRNAs, their strong secondary structure, and the high frequency of post-transcriptional tRNA modifications, high-throughput experimental quantification of tRNA expression levels is challenging. Two dedicated experimental high-throughput approaches for the quantification of tRNA expression, hydro-tRNA-seq (Gogakos et al., 2017) and DM-tRNA-seq (Zheng et al., 2015), have been proposed and were applied in human HEK293 cells. smallRNA-seq and H3K27ac-ChIP-seq data were also used previously to quantify tRNA expression (Gingold et al., 2014; Hernandez-Alias et al., 2020; Ji et al., 2015), and these data are more widely available for different human cell types. We compared relative tRNA expression levels based on these methods for HEK293 with those from hydro-tRNA-seq and DM-tRNA-seq (Figure S7A), and found that smallRNA-seq-based relative tRNA abundances correlated better with those from hydro-tRNA-seq and DM-tRNA-seq data. As a confirmation, HEK-specific tAIs calculated using smallRNA-seq-based tRNA abundances showed the most significant difference between expressed and highly expressed cytoplasmic mRNAs, while tAIs calculated using H3K27ac-ChIP-seq-based tRNA abundances showed no difference between both (Figure S7B). Increased tAIs for mRNAs with highest cytoplasmic expression would be expected, based on previous analyses of mRNA codon biases. Thus, we used smallRNA-seq-based tRNA abundances to calculate cell type-specific tAIs for all cell lines.

In the following, the quantification of tRNA abundances based on different experimental data is explained. Effective tRNA abundances are calculated from these as described above.

#### (a) based on smallRNA-seq data

Fastq files with smallRNA-seq reads were downloaded for different species and cell types from various sources (see Table S3). Reads were pre-processed using the fastx toolkit (http://hannonlab.cshl.edu/fastx_toolkit/) and then mapped to native and mature tRNA sequences using segemehl v0.2 (Hoffmann et al., 2009). Of the mapped reads, only those with a minimum length of 15 nucleotides were retained. To account for the high frequency of tRNA modifications, which may result in mapping mismatches, the allowed mismatch ratio (mismatched nucleotides / read length) was set to <=10%. (Other mismatch ratio cutoffs, <7% and <15%, were also tested, but did not improve the correlation with hydro-tRNA-seq (Gogakos et al., 2017) and DM-tRNA-seq (Zheng et al., 2015), or resulted in a smaller fraction of reads mapping to tRNA sequences in sense direction; data not shown). tRNA abundances were calculated as the number of smallRNA-seq reads mapping to each tRNA anticodon type.

#### (b) based on H3K27ac ChIP-seq data for HEK293

Bedfiles of H3K27ac-ChIP-seq peak regions identified in HEK293 cells were downloaded from ENCODE (www.encodeproject.org) (Consortium and The ENCODE Project Consortium, 2004) (see Table S3). H3K27ac ChIP-seq peak regions were overlapped with tRNA genes (extended by 500 nucleotides up- and downstream) using bedtools (Quinlan, 2014). tRNA abundances were calculated for each tRNA anticodon type as the sum of peak enrichment values (given in column 7 of the bedfiles as log2 fold enrichment H3K27ac-ChIP-seq over background) of peaks overlapping extended tRNA gene annotations.

#### (c) based on hydro-tRNA-seq and DM-tRNA-seq in HEK293

Hydro-tRNA-seq-based tRNA quantifications were taken from Table S5 of Gogakos et al. (Gogakos et al., 2017). DM-tRNA-seq (Zheng et al., 2015) read counts of two replicates were downloaded from GEO (www.ncbi.nlm.nih.gov/geo, accession numbers GSM1624820 and GSM1624821) and tRNA abundances were calculated as the average of the two replicates.

### tRNA adaptation index (tAI)

As proposed by dos Reis et al. (dos Reis et al., 2004), the tAI of an ORF was calculated as the geometric mean of normalized effective tRNA abundances complementary to codons in the ORF:

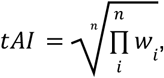

where *n* is the total number of codons in an ORF, and *w_i_* is the normalized effective tRNA abundance of the tRNA anticodon complementary to the codon at position*i*.

Normalized tRNA abundances were obtained by dividing each effective tRNA abundance by the maximum of all effective tRNA abundances:

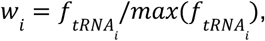

where 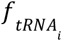 is the frequency of the tRNA complementary to the codon at position*i*.

We chose to normalize to the maximum of all effective tRNA abundances, as opposed to the maximum among synonymous codons coding for the same amino acid, because we wanted to analyse the global correspondence between tRNA abundances and codon usage, independent of amino acid identities. For comparison, synonymous-tAIs calculated considering relative tRNA abundances for synonymous codon usage are shown in Figure S5B.

First-10-codon, first-20-codon or first-30-codon tAIs were calculated by considering only the first 10, 20 or 30, respectively, codons downstream of AUG start codons.

### Randomized control sequences

Shuffled sequences were generated by randomly permuting nucleotides in an ORF. Shuffled tAIs were calculated for codon frequencies from 100 random permutations.

Frame-shifted tAIs were calculated from codon frequencies in sequences starting one and two nucleotides downstream of start codons and ending two and one, respectively, nucleotides upstream of stop codons of ORFs.

### Adaptation of the correlation test for translational selection from dos Reis et al. (2004)

To test for translational selection on mRNA codon usage in a species, dos Reis et al. (dos Reis et al., 2004) proposed a correlation test, in particular the correlation between tAI and the effective number of codons, adjusted for the GC bias at the third codon position of mRNAs. Here, we adapted this translational selection test to measure the global correspondence between codon usage bias and tRNA abundances for all 60 codons (excluding start and stop codons), not just the correspondence within groups of synonymous codons. For that, we calculated the effective number of codons independent of amino acid identities and adjusted for the GC bias at all three codon positions. Confidence intervals of correlation coefficients in Figure 3C were estimated by 1000 times bootstrapping. All calculations were implemented in python.

### Quantification of cytoplasmic gene expression levels per species

In Figures 3, S5 and S6, we sought to estimate cytoplasmic RNA expression levels for three species (human, fly and fission yeast) by combining cytosolic gene expression quantifications from different studies (see Table S3 for sources and accession codes). For human, we used data from five cell lines, GM12878, HeLa-S3, HEK293, HepG2 and K562 (Consortium and The ENCODE Project Consortium, 2004; Subtelny et al., 2014), for fly we used data from early embryos and two cell lines, S2 and ML-DmD17-c3 (Aspden et al., 2014; Bouvrette et al., 2018; Li et al., 2016), and for fission yeast we used data from two different growth conditions (Herzel et al., 2018; Subtelny et al., 2014). For four human cell lines, except HEK293, we downloaded gene quantifications from ENCODE (www.encodeproject.org). For HEK293 and other species, we downloaded fastq files with polyA-selected RNA-seq reads obtained from cytosolic RNA fractions, preprocessed them using the fastx toolkit (http://hannonlab.cshl.edu/fastx_toolkit/), and quantified gene expression levels using RSEM (Li and Dewey, 2011) with STAR (Dobin et al., 2013). Gene expression levels were averaged over replicates. To combine expression data from different experiments for each species, we first ranked genes by their cytoplasmic expression level in each experiment. We then used the maximum rank across experiments as the combined rank for each gene. Combined ranks were normalized to the maximum combined rank, so that genes with highest cytosolic expression levels had combined ranks close to 1. Top cytoplasmic lincRNAs were those with a combined rank > 0.85, and top cytoplasmic mRNAs were those with a combined rank > 0.99.

### Quantification of total and cytosolic transcript expression levels in human cell lines

In Figures 4, 5, S7, S8 and S9, we used total and cytoplasmic transcript expression levels for five human cell lines (GM12878, HeLa-S3, HEK293, HepG2 and K562; see Table S3 for sources and accession codes). Transcript quantifications for four cell lines, except HEK293, were downloaded from ENCODE (Consortium and The ENCODE Project Consortium, 2004). For HEK293, polyA-selected RNA-seq reads (Aktaş et al., 2017; Subtelny et al., 2014) were downloaded and preprocessed using the fastx toolkit (http://hannonlab.cshl.edu/fastx_toolkit/), and transcript expression levels were quantified using RSEM (Li and Dewey, 2011) with STAR (Dobin et al., 2013). Expression levels were averaged over replicates. RNAs with TPM > 0.1 were considered expressed. The threshold for defining cytoplasmic RNAs was set to the first-quartile (25%) of the mRNA cytosolic expression values, for each cell type. We excluded histone mRNAs, as these are not usually polyadenylated and result in incorrect expression values in polyA-selected RNA-Seq.

In Figure 4B, mRNAs with cytoplasmic expression levels matching those of lincRNAs were selected as follows. First, for each cytoplasmic lincRNA all mRNAs were identified that had similar cytoplasmic expression levels (identical values after log10-transformation and rounding to 2 decimal places). From these mRNAs, 10 were randomly sampled with replacement for each cytoplasmic lincRNA.

### Quantification and analysis of Ribo-Seq data in human cell lines

Ribo-Seq data for five human cell lines were downloaded from several studies ((Cenik et al., 2015; Huang et al., 2019; Martinez et al., 2020; Solomon et al., 2017; Subtelny et al., 2014); see Table S3). Adapter sequences were trimmed from read ends using cutadapt v1.8 (Martin, 2011), and reads were retained with a length between 16 to 35 nucleotides and a quality score of >=30 in at least 90% of the reads’ bases. Reads were discarded that mapped to human rRNAs or tRNAs (ENSEMBL database v91 (Zerbino et al., 2018)) using bowtie2 v2.3.0 (−L 15 −k 20) (Langmead and Salzberg, 2012). Reads were further discarded that mapped to two or more mRNA coding regions or longest lincRNA ORFs.

The position of the ribosome P site within Ribo-Seq reads was determined from reads overlapping mRNA start codons. In particular, the three most frequent distances of the AUG start codon from the read start (read offset) were considered for each read length (if they were found in more than 500 reads and in more than 1% of reads overlapping mRNA start codons). Then, for each longest mRNA coding region or longest lincRNA ORF, Ribo-Seq reads were counted if the corresponding ribosome P site was in-frame considering the three read offsets for that respective Ribo-Seq read length.

Relative ribosome-binding was calculated for each longest coding region/ORF as the log2 ratio of the normalized Ribo-Seq read count (plus a pseudo-read count of 0.1) to the cytosolic expression level (FPKM) of the transcript harboring the longest coding region/ORF. Normalized Ribo-Seq read counts were calculated by dividing by the length of coding region/ORF and the total number Ribo-Seq reads mapping in-frame to mRNA coding regions, and by multiplying by 1e+9.

We excluded from this analysis histone mRNAs, as these are not usually polyadenylated and result in incorrect expression values in polyA-selected RNA-Seq, and snoRNAs, as these may associate with ribosomes that translate other RNAs.

## Acknowledgements

We would like to thank the ENCODE consortium and other experimental labs for generating the publicly available data used in this study. This work was supported by the Swiss National Science Foundation (grant FN 310030_152724/1).

## Author Contributions

AB, ACM and SB designed the project. AB carried out the computational analyses and prepared the results. RD mapped Ribo-Seq data. AB, ACM, and SB discussed results and wrote the paper.

---

**Figure S1:**
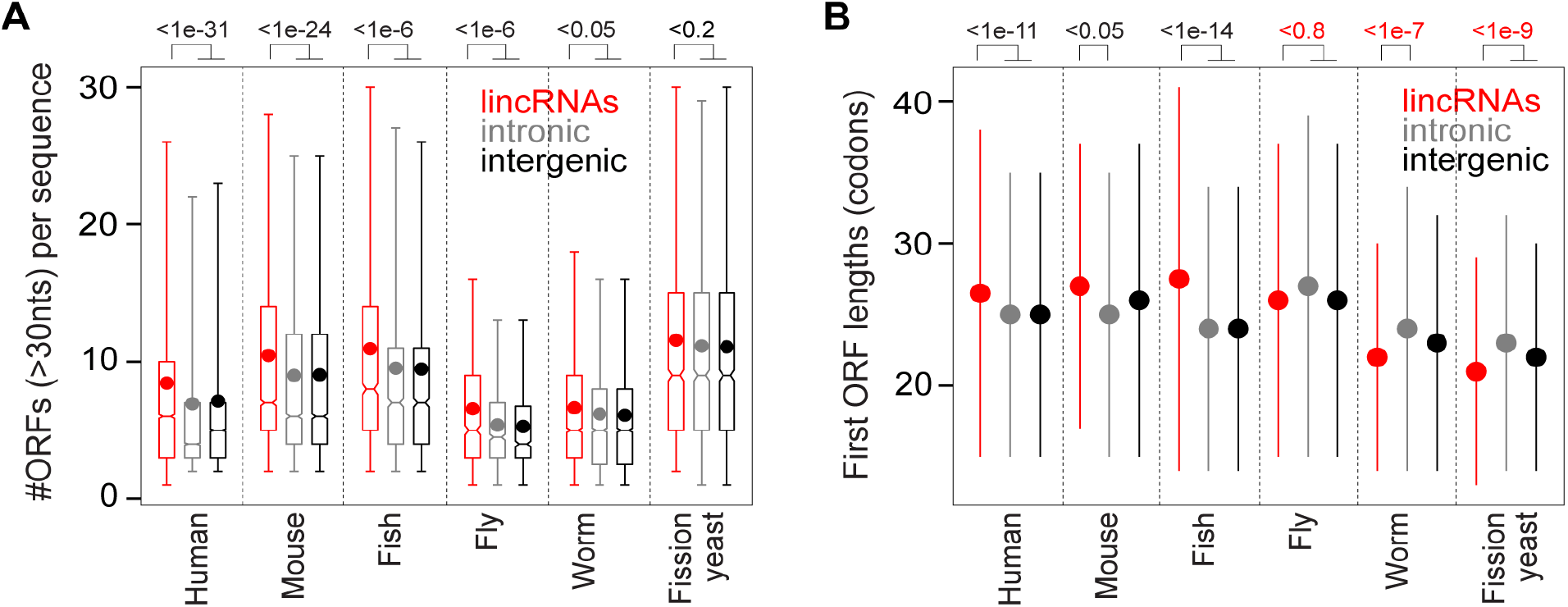
Additional ORF identification statistics (related to Figure 1A and B) (A) Boxplots of number of ORFs (>10 codons) in lincRNAs (red) and in intronic (gray) and intergenic (black) control regions for different species. Dots indicate the average number of ORFs per sequence. P values are from Wilcoxon’s Rank sum test. (B) Median lengths of first ORFs in lincRNAs (red), and in size-matched and G+C content-matched intronic (gray) and intergenic (black) control regions. Error bars represent median absolute deviations. P values are indicated from Wilcoxon’s rank-sum test. P values are marked red, if the median lincRNA value is below the value for control regions.

**Figure S2:**
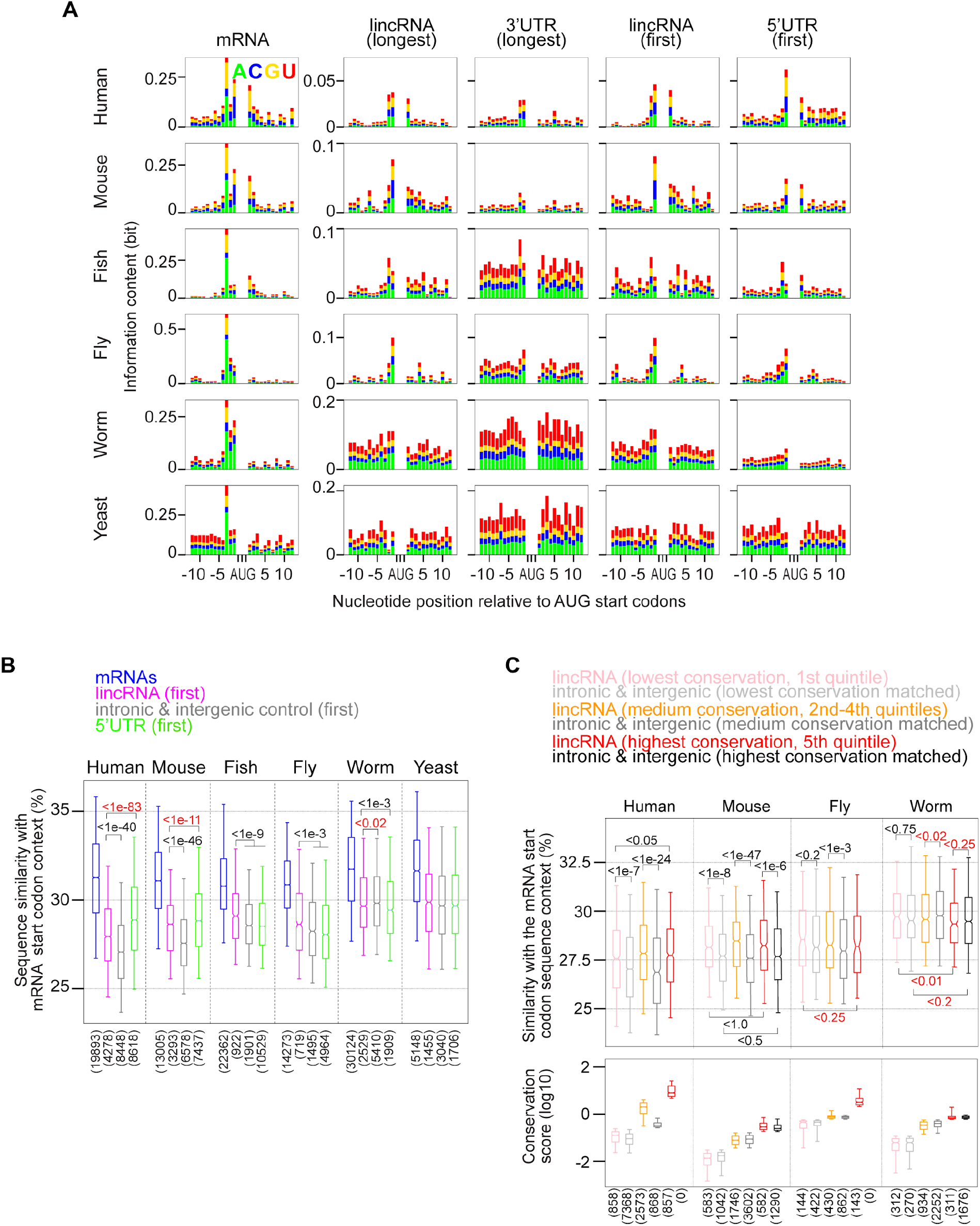
Start codon sequence context (related to Figure 1C) (A) Information content (see Methods) for the region +/−12 nucleotides around AUG start codons for different ORFs (columns, indicated on top) and species (rows, indicated left). The sequence motif around mRNA start codons shows some similarity with the Kozak consensus sequence (gcc(A/G)cc<u>AUG</u>G). (B) Sequence similarity with the consensus sequence motif of the region +/−12 nucleotides around mRNA start codons (see Methods) for mRNAs (blue), lincRNA first ORFs (pink), intronic and intergenic control ORFs (gray), and first ORFs in 5’UTRs (light green). P values are indicated from Wilcoxon’s rank-sum test. (C) Sequence similarity with the consensus sequence motif of the region +/−12 nucleotides around mRNA start codons for longest ORFs in lincRNAs that have low (1st quintile, pink), intermediate (2nd to 4th quintile, orange) or high conservation scores (5th quintile, red) (see Methods). As a comparison, the sequence similarity is shown for longest ORFs in intronic and intergenic control regions with conservation scores corresponding to those of lincRNAs (i.e. within lincRNA quintile cuts; color code indicated above). The distributions of conservation scores are shown as boxplots in the lower panel. Conservation data was not available for fish and fission yeast. P values are indicated from Wilcoxon’s rank-sum test. P values are marked red, if the median lincRNA value is below the value for control regions.

**Figure S3:**
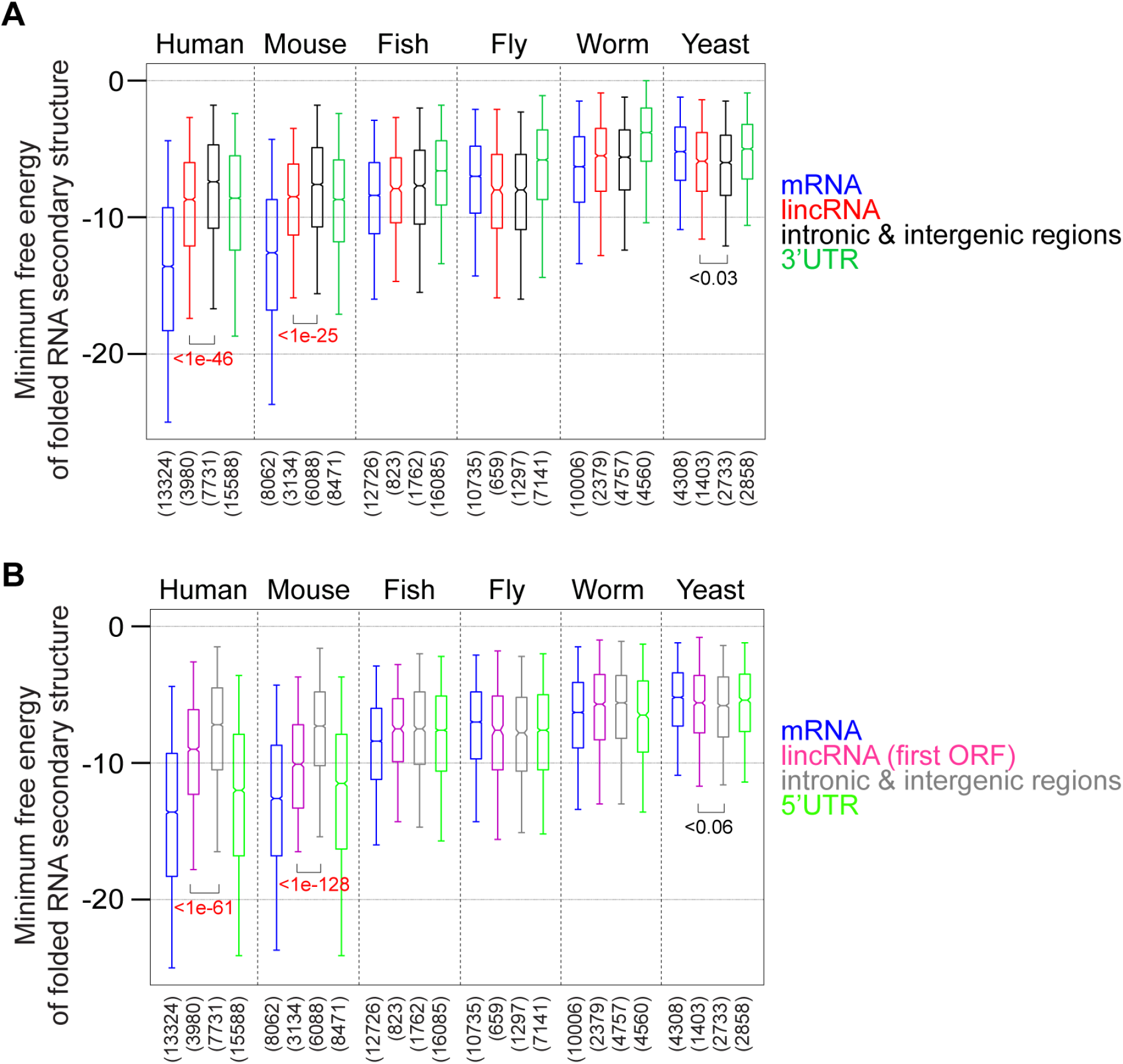
Structure context around AUG start codons in different RNA types and species. (A, B) Minimum free energy (MFE) of RNA structures folded from the region +/− 25 nucleotides around AUG start codons in different species (indicated on top) and RNA types (color coded with labels on the right of each panel). P values are indicated from Wilcoxon’s rank-sum test to compare MFEs for lincRNAs with those of control ORFs. P values are marked red, if lincRNA median values are below those of intronic and intergenic controls. Only significant p values (p<0.05) are indicated.

**Figure S4:**
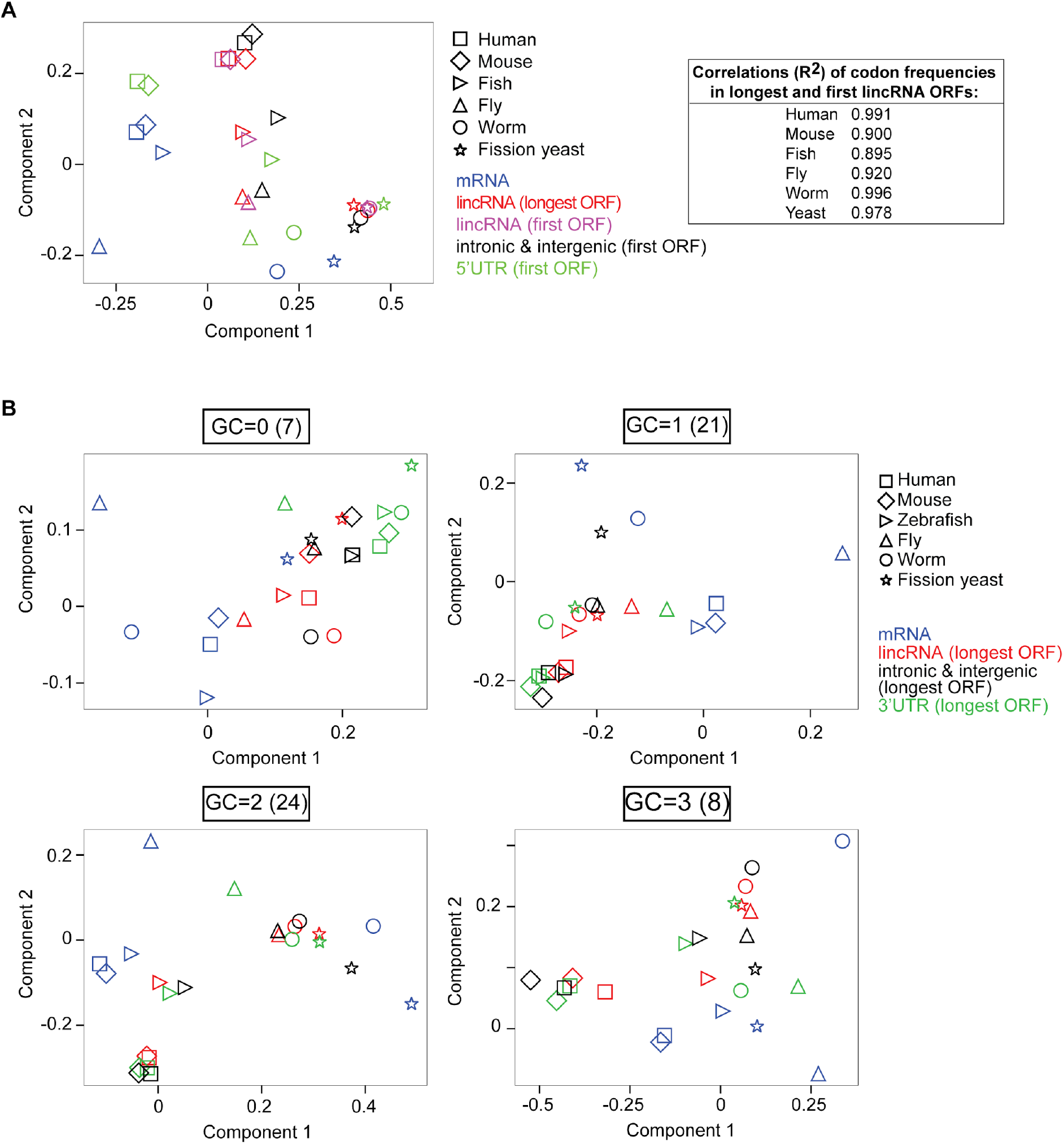
Multiple correspondence analysis of codon counts for additional ORFs and for codons stratified by G+C content. (A) Left panel: First two components from a multiple correspondence analysis performed on trinucleotide (“codon”) counts (excluding start and stop codons) in mRNA coding regions (blue), lincRNA longest ORFs (red), lincRNA first ORFs (pink), first ORFs in intronic and intergenic control ORFs (black) and first ORFs in 5’UTRs (light green) for six species (represented by different symbols). Right panel: Squared correlation coefficient (r^2^) between codon frequencies in longest and first ORFs of lincRNAs. (B) Component 1 versus 2 plot from a multiple correspondence analysis carried out with counts of codons with different number of G and C nucleotides, as indicated above each panel, for mRNA coding regions (blue), lincRNA longest ORFs (red), longest ORFs in intronic and intergenic control regions (black) and longest ORFs in 3’UTRs (green) for six species (represented with different symbols). The number of codons with a certain number of G and C nucleotides is indicated in parenthesis above each panel.

**Figure S5:**
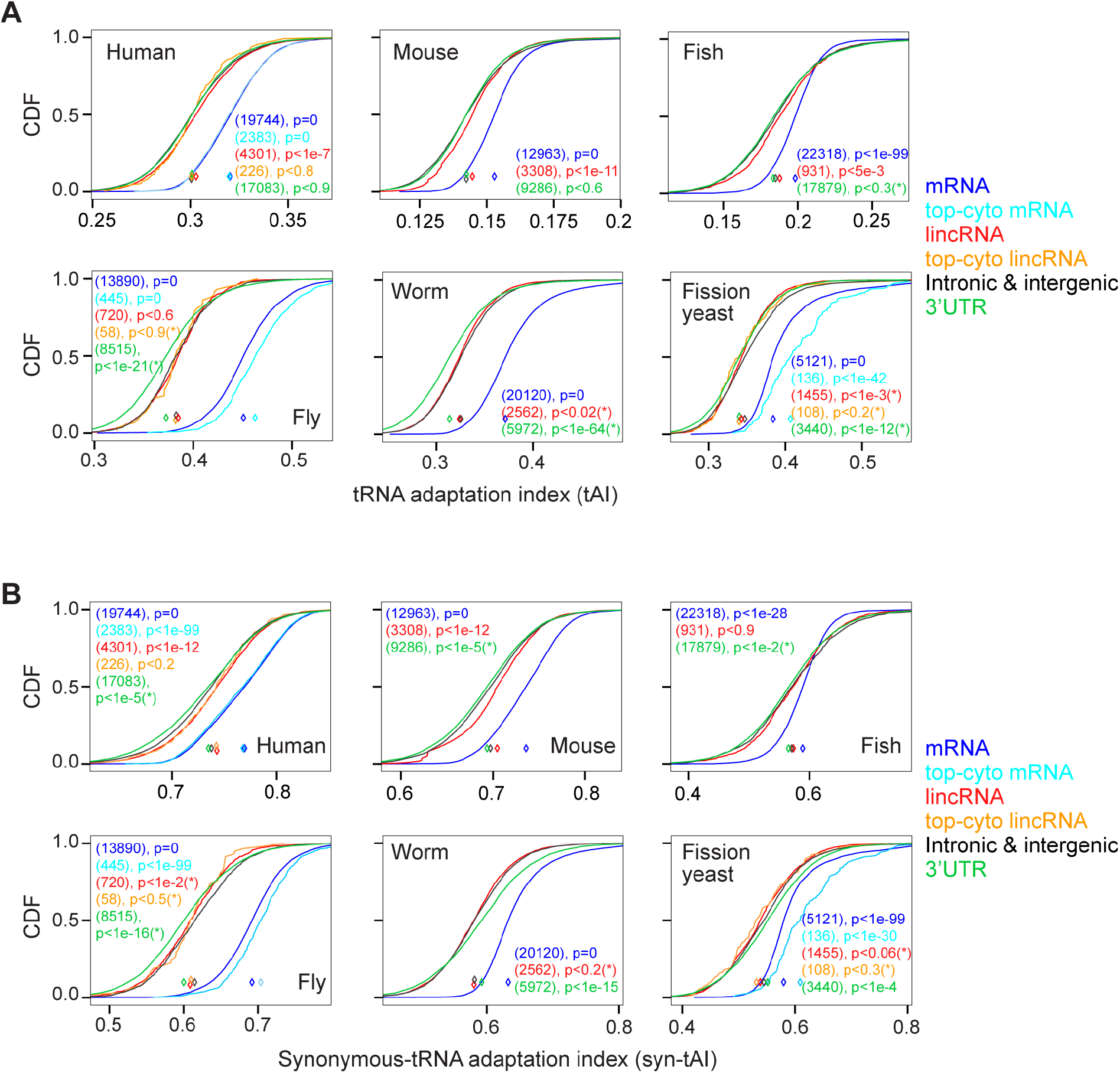
Cumulative distributions of tRNA adaptation indexes (tAIs) for different RNA types and species. (A) tRNA adaptation index (tAI, calculated using tRNA gene counts; see Methods) for mRNA coding regions (blue), and longest ORFs in lincRNAs (red), intronic and intergenic control regions (black) and 3’UTRs (green) in different species (see label inside each panel). In the case of human, fly, and yeast, tAIs of top-cytoplasmic mRNAs (light blue) and cytoplasmic lincRNAs (orange) are shown in addition. P values are indicated from Wilcoxon’s rank-sum test comparing tAIs of control ORFs with those of ORFs in other RNA types (color-coded). P values are marked with an asterisk if the median tAI is below the one of control ORFs. (B) Synonymous-tAI, quantifying the correspondence between synonymous codon usage and relative abundances of tRNAs decoding the same amino acid, for the ORFs types and in all species as in (A).

**Figure S6:**
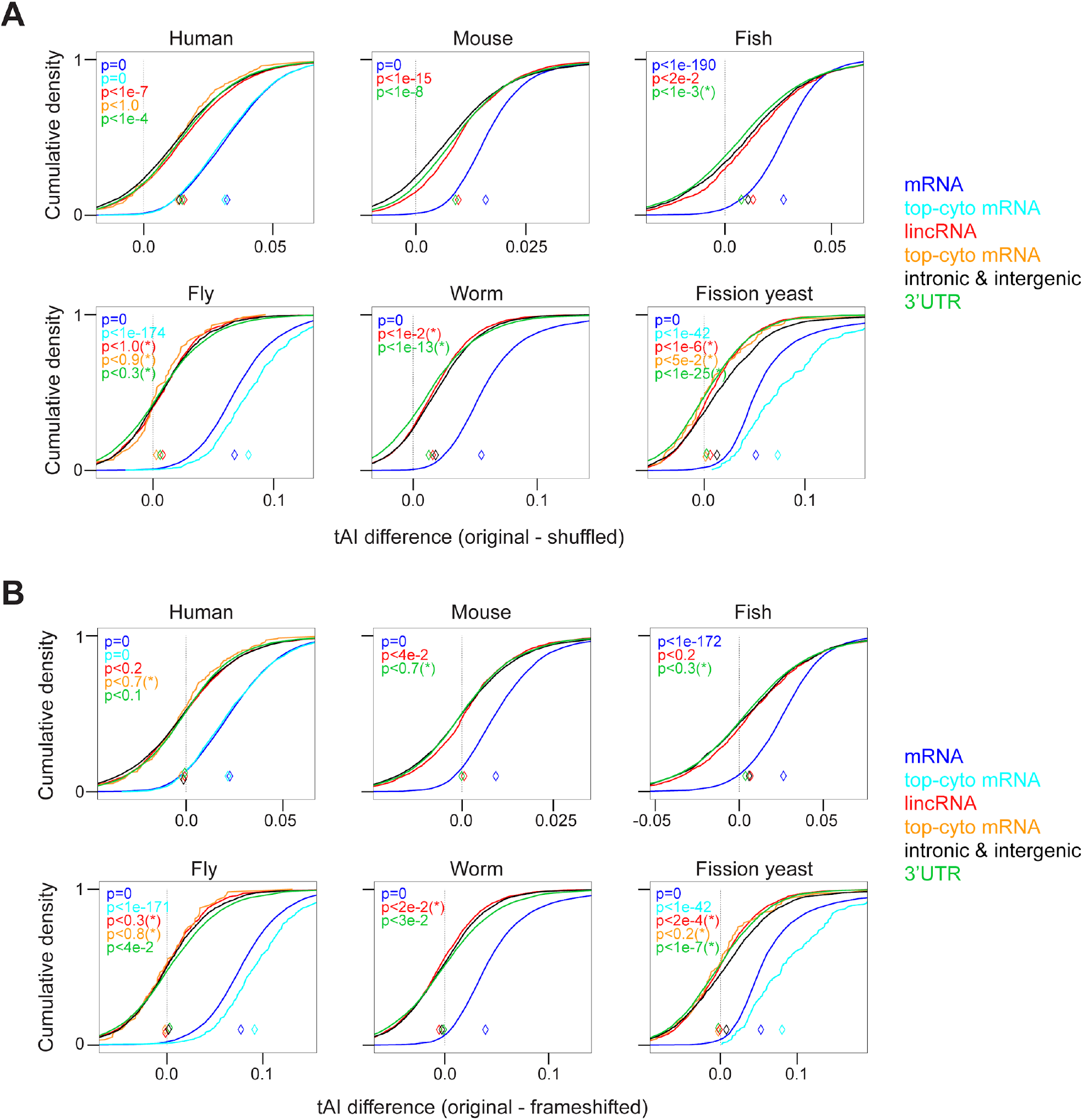
Comparison of tAIs between original and shuffled or frame-shifted ORFs in six species. Cumulative distributions of tAI differences between (A) original and shuffled ORFs, and (B) original and frame-shifted ORFs for six species (indicated on top of each panel). tAI differences were calculated for mRNA coding regions (blue), longest ORFs in lincRNAs (red), intronic and intergenic control regions (black), and 3’UTRs (green). For human, fly, and yeast, tAI differences of top cytoplasmic mRNAs (light blue) and cytoplasmic lincRNAs (orange) are shown in addition. P values were calculated using Wilcoxon’s rank sum test to compare the distribution of tAI differences for control regions with those of other RNA types (color-coded). P values are marked with asterisks if the median tAI difference of that set of ORFs is below that of control ORFs.

**Figure S7:**
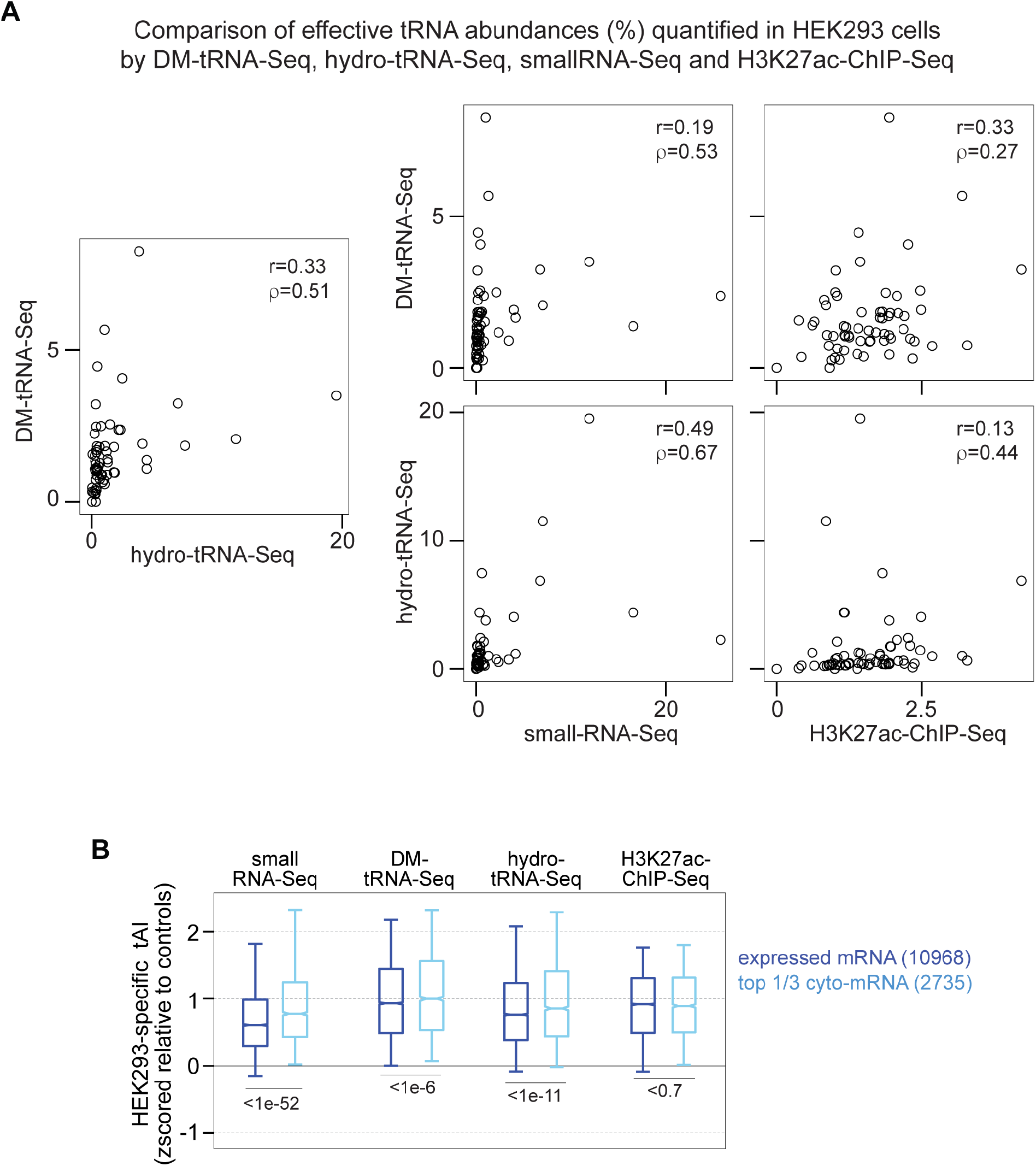
Comparison of tRNA abundances and cell type-specific tAIs derived from different experimental approaches in HEK293 cells. (A) Left panel: Comparison of tRNA anticodon frequencies derived from two dedicated experimental approaches, hydro-tRNA-Seq (Gogakos et al. 2017; x-axis) and DM-tRNA-Seq (Zheng et al. 2015; y-axis). Pearson (r) and Spearman (ρ) correlation coefficients are indicated. Right four panels: Comparison between tRNA anticodon frequencies derived from one of the two dedicated experimental approaches (see y-axis labels) with those from smallRNA-Seq or H3K27ac-ChIP-Seq (see x-axis label; see Methods). Pearson (r) and Spearman (ρ) correlation coefficients are indicated. (B) Box plots of cell type-specific tAIs (z-scored to cell type-specific tAIs of controls) in HEK293 cells calculated using effective tRNA abundances quantified from different experimental approaches, as indicated on top. For each experimental approach, cell type-specific tAIs are shown for expressed mRNAs (blue) and for top 1/3 cytoplasmic mRNAs (light blue). The number of ORFs in each group are indicated in parenthesis on the right. P values are calculated using Wilcoxon’s rank-sum test. Cell type-specific tAIs in HEK293 show the most significant difference between expressed and top cytoplasmic mRNAs when using smallRNA-Seq-derived tRNA abundances.

**Figure S8:**
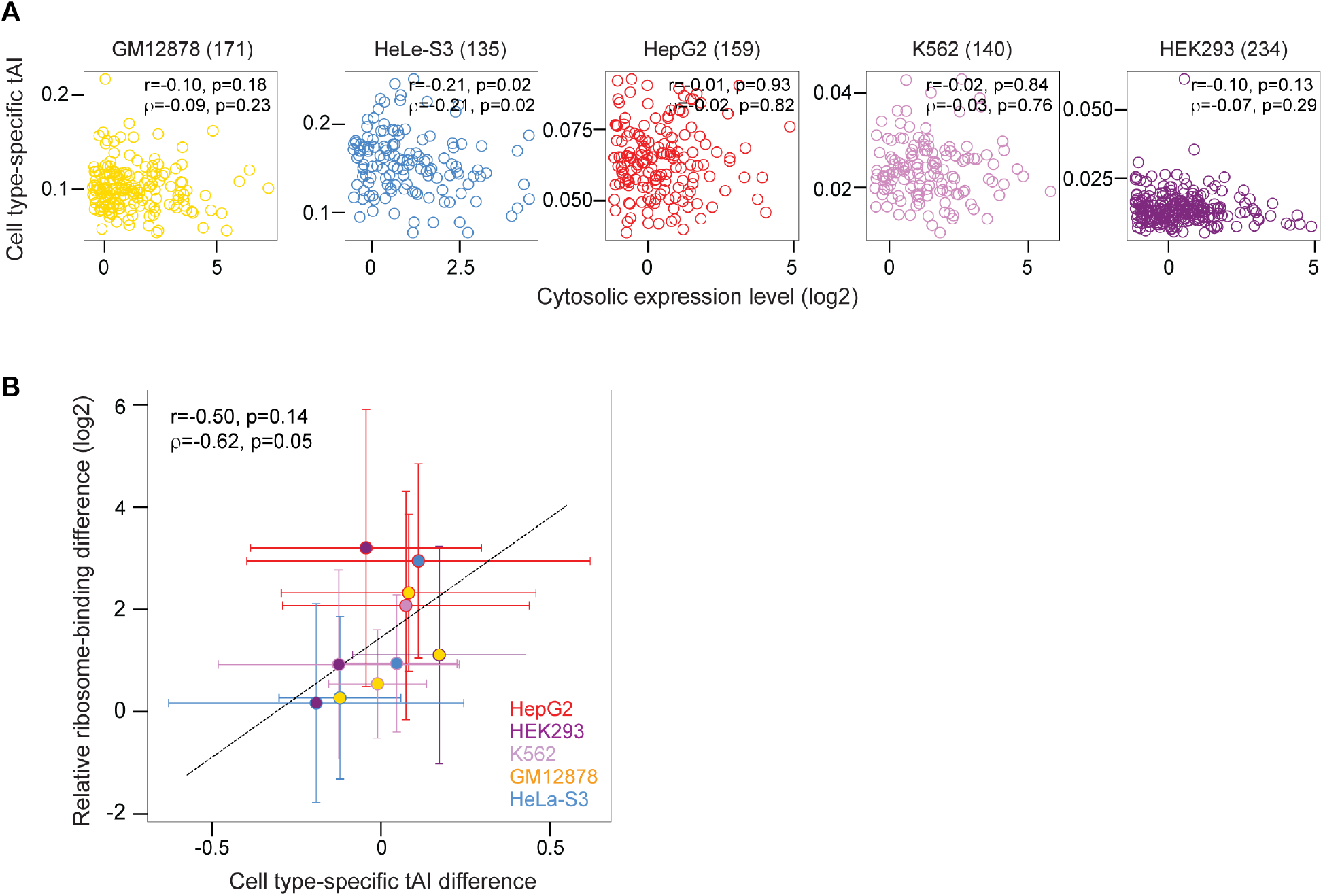
Relationship between expression level and cell type-specific tAI and ribosome-binding for cytoplasmic lincRNAs in five human cell lines (related to Figure 4) (A) Scatter plot of cytoplasmic expression levels and tAIs of cytoplasmic lincRNAs in five human cell lines (indicated on top of each panel with number of cytoplasmic lincRNAs in parenthesis). Pearson (r) and Spearman (ρ) correlation coefficients are indicated with p values. (B) Scatter plot of median differences in cell type-specific tAIs (z-scored to tAIs of control regions) and median differences in relative ribosome-binding for lincRNAs that are cytoplasmic in a pair of cell lines (color code on the right with number of cytoplasmic lincRNAs shared between two cell lines in parenthesis). Error bars represent the median absolute deviation. Pearson (r) and Spearman (ρ) correlation coefficients between median values are indicated with p values.

**Figure S9:**
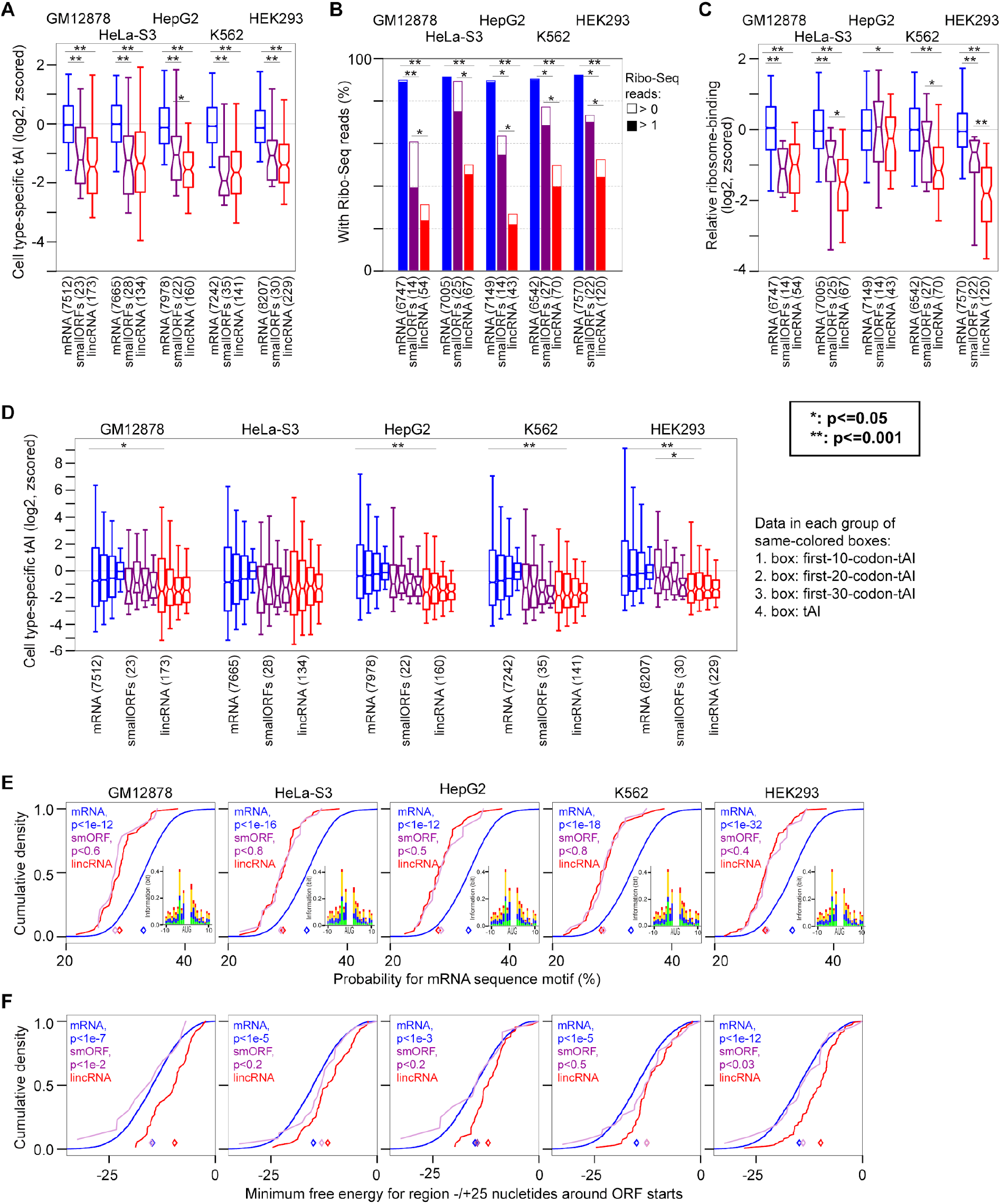
Comparisons of tAIs and ribosome-binding for mRNAs, small protein-encoding RNAs and lincRNAs in five human cell lines. (A) Boxplot of cell type-specific tAIs (log2-transformed and z-scored relative to tAIs of cytoplasmic mRNA coding regions) for mRNA coding regions (blue), small protein-encoding ORFs (smORFs; purple) and lincRNA longest ORFs (red) of cytosolic RNAs in five human cell lines (indicated on top). Number of RNAs in each box is indicated below each box. Asterisks indicate p values from Wilcoxon’s rank-sum test (*:<=0.05 and **:<=0.001). (B) Fraction of ORFs with in-frame Ribo-Seq reads (filled bars with more than one Ribo-Seq read and empty bars with one Ribo-Seq read). The number of ORFs with Ribo-Seq reads is indicated below each bar. Asterisks represent p values from Fisher’s exact test (*:<=0.05 and **:<=0.001). (C) Boxplot of relative ribosome-binding (calculated as the log2-ratio of normed Ribo-Seq reads to cytoplasmic expression; z-scored relative to that of mRNA coding regions) for the same types of cytoplasmic RNAs as in (A) with Ribo-Seq reads. Cell lines are indicated above. Number of RNAs included in each box is indicated below each box. Asterisks represent p values from Wilcoxon’s rank-sum test (*:<=0.05 and **:<=0.001). (D) Cell type-specific tAIs (log2-transformed and z-scored as in (A)) for different regions within ORFs in cytoplasmic RNAs: tAI for the first 10, 20, and 30 codons downstream of start codons are shown in first, second, and third box, respectively; tAI of entire ORFs (fourth box) for each RNA type and in each cell line. Asterisks represent p values from Wilcoxon’s rank-sum test (*:<=0.05 and **:<=0.001). (E) Sequence context around start codons of three types of cytoplasmic RNAs: mRNA (blue), smORFs (purple), and lincRNAs (red). Each panel shows the cumulative distribution of the probabilities for the consensus mRNA sequence motif of cytoplasmic RNAs in a human cell line. P values are calculated using Wilcoxon’s rank-sum test to compare lincRNAs with the two other types of RNA. (F) RNA structure context around start codons of the three types of cytoplasmic RNAs. Each panel shows the cumulative distribution of the minimum free energy of the region −/+ 25 nucleotides around start codons for cytoplasmic RNAs in a human cell line. P values are calculated using Wilcoxon’s rank-sum test to compare lincRNAs with the two other types of RNA.

**Table S1:** Number of analyzed mRNA coding regions and lincRNAs with longest ORF > 10 codons in different species. The final number of lincRNAs analysed in each species is marked bold.

|  | **Human** | **Mouse** | **Fish** | **Fly** | **Worm** | **Fission yeast** |
| --- | --- | --- | --- | --- | --- | --- |
| Number of mRNAs coding regions | 19744 | 12963 | 22318 | 13890 | 20120 | 5121 |
| Number of annotated lincRNAs (isoform with longest ORF) | 4867 | 3929 | 1203 | 1262 | 3242 | **1455** |
| lincRNAs without PhyloCSF novel track overlap | 4695 | 3791 | - | 1078 | 3011 | - |
| lincRNAs without simple repeat or low complexity overlap | 4370 | **3308** | **931** | **720** | **2542** | - |
| lincRNAs without peptide-encoding ORF overlap | **4301** | - | - | - | - | - |

**Table S2:**
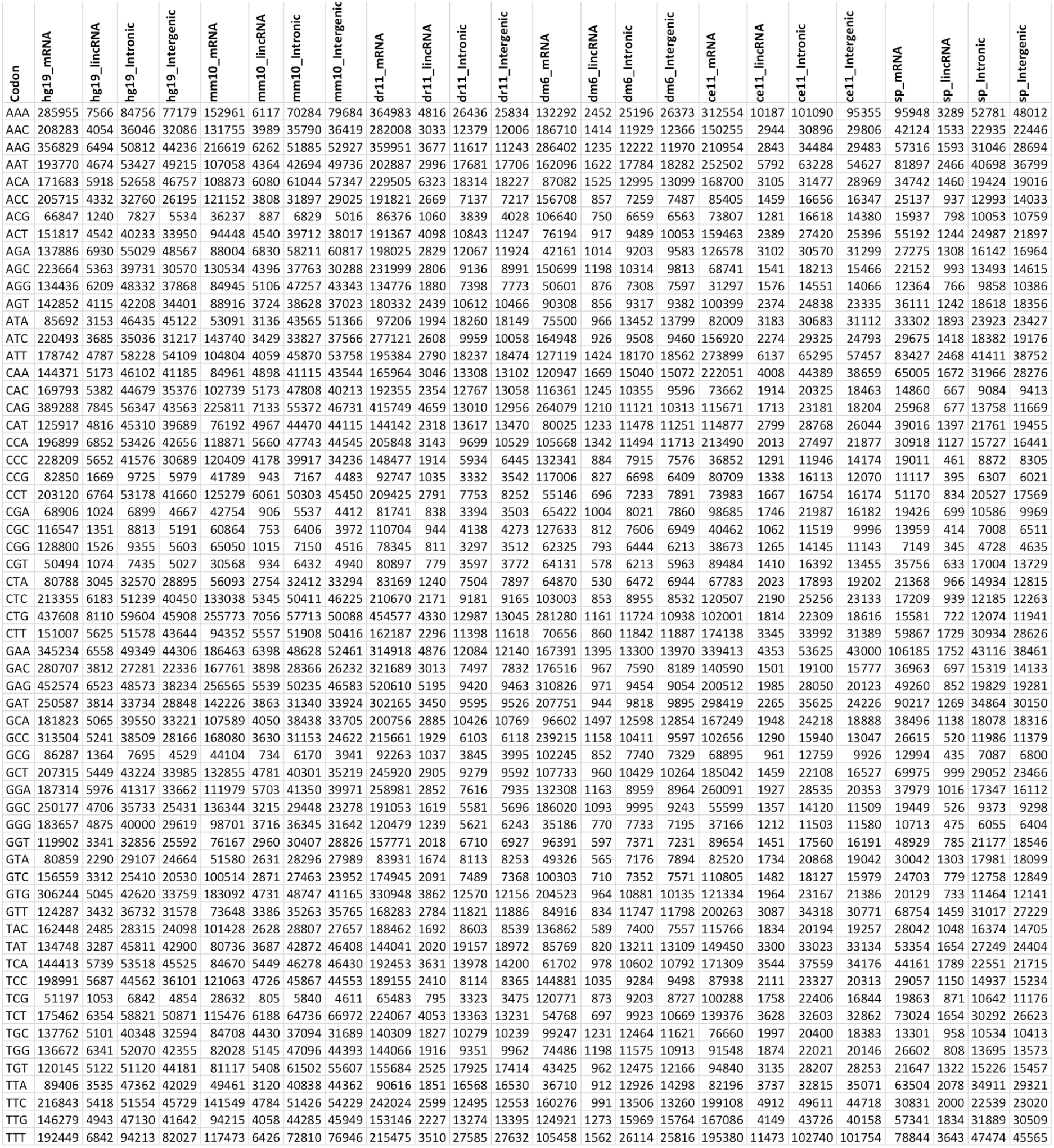
Codon counts for ORFs in different RNA types and species.

**Table S3:** Sources and accession codes of experimental data used in this study.

| Organism | Cell type | Experimental assay | Source | Accessions | Reference |
| --- | --- | --- | --- | --- | --- |
| Human | GM12878 | smallRNA-Seq | www.encodeproject.org | ENCSR000CRE | The ENCODE Project Consortium, The ENCODE (ENCyclopedia Of DNA Elements) Project. Science 306, 636–640 (2004) |
| Human | GM12878 | total polyA-RNA-Seq | www.encodeproject.org | ENCSR000COQ | The ENCODE Project Consortium, The ENCODE (ENCyclopedia Of DNA Elements) Project. Science 306, 636–640 (2004) |
| Human | GM12878 | cytosolic polyA-RNA-Seq | www.encodeproject.org | ENCSR000COR | The ENCODE Project Consortium, The ENCODE (ENCyclopedia Of DNA Elements) Project. Science 306, 636–640 (2004) |
| Human | GM12878 | Ribo-Seq | www.ncbi.nlm.nih.gov/geo/ | GSM1609378, GSM1609379 | C. Cenik, et al., Integrative analysis of RNA, translation, and protein levels reveals distinct regulatory variation across humans. Genome Res. 25, 1610–1621 (2015) |
| Human | HeLa-S3 | smallRNA-Seq | www.encodeproject.org | ENCSR000CRQ | The ENCODE Project Consortium, The ENCODE (ENCyclopedia Of DNA Elements) Project. Science 306, 636–640 (2004) |
| Human | HeLa-S3 | total polyA-RNA-Seq | www.encodeproject.org | ENCSR000CPR | The ENCODE Project Consortium, The ENCODE (ENCyclopedia Of DNA Elements) Project. Science 306, 636–640 (2004) |
| Human | HeLa-S3 | cytosolic polyA-RNA-Seq | www.encodeproject.org | ENCSR000CPP | The ENCODE Project Consortium, The ENCODE (ENCyclopedia Of DNA Elements) Project. Science 306, 636–640 (2004) |
| Human | HeLa-S3 | Ribo-Seq | www.ncbi.nlm.nih.gov/geo/ | GSM3566406, GSM3566407, GSM3566408, GSM3566409 | T. F. Martinez, et al., Accurate annotation of human protein-coding small open reading frames. Nature Chemical Biology 16, 458–468 (2020) |
| Human | HepG2 | smallRNA-Seq | www.encodeproject.org | ENCSR000CRX | The ENCODE Project Consortium, The ENCODE (ENCyclopedia Of DNA Elements) Project. Science 306, 636–640 (2004) |
| Human | HepG2 | total polyA-RNA-Seq | www.encodeproject.org | ENCSR000CPE | The ENCODE Project Consortium, The ENCODE (ENCyclopedia Of DNA Elements) Project. Science 306, 636–640 (2004) |
| Human | HepG2 | cytosolic polyA-RNA-Seq | www.encodeproject.org | ENCSR000CPF | The ENCODE Project Consortium, The ENCODE (ENCyclopedia Of DNA Elements) Project. Science 306, 636–640 (2004) |
| Human | HepG2 | Ribo-Seq | www.ncbi.nlm.nih.gov/geo/ | GSM2676361, GSM2676363 | O. Solomon, et al., RNA editing by ADAR1 leads to context-dependent transcriptome-wide changes in RNA secondary structure. Nat. Commun. 8, 1440 (2017) |
| Human | HepG2 | Ribo-Seq | www.ncbi.nlm.nih.gov/geo/ | GSM3450419, GSM3450420 | H. Huang, et al., Histone H3 trimethylation at lysine 36 guides m6A RNA modification co-transcriptionally. Nature 567, 414–419 (2019) |
| Human | K562 | smallRNA-Seq | www.encodeproject.org | ENCSR000CRF | The ENCODE Project Consortium, The ENCODE (ENCyclopedia Of DNA Elements) Project. Science 306, 636–640 (2004) |
| Human | K562 | total polyA-RNA-Seq | www.encodeproject.org | ENCSR000CPH | The ENCODE Project Consortium, The ENCODE (ENCyclopedia Of DNA Elements) Project. Science 306, 636–640 (2004) |
| Human | K562 | cytosolic polyA-RNA-Seq | www.encodeproject.org | ENCSR000COK | The ENCODE Project Consortium, The ENCODE (ENCyclopedia Of DNA Elements) Project. Science 306, 636–640 (2004) |
| Human | K562 | Ribo-Seq | www.ncbi.nlm.nih.gov/geo/ | GSM3566410, GSM3566411, GSM3566412 | T. F. Martinez, et al., Accurate annotation of human protein-coding small open reading frames. Nature Chemical Biology 16, 458–468 (2020) |
| Human | HEK293 | smallRNA-Seq | www.ncbi.nlm.nih.gov/geo/ | GSM1067868 | S. Kishore, et al., Insights into snoRNA biogenesis and processing from PAR-CLIP of snoRNA core proteins and small RNA sequencing. Genome Biol. 14, R45 (2013) |
| Human | HEK293 | total polyA-RNA-Seq | www.ncbi.nlm.nih.gov/geo/ | GSM2258993, GSM2258994, GSM2258996 | T. Aktaş, et al., DHX9 suppresses RNA processing defects originating from the Alu invasion of the human genome. Nature 544, 115–119 (2017) |
| Human | HEK293 | cytosolic polyA-RNA-Seq | www.ncbi.nlm.nih.gov/geo/ | GSM1276541 | A. O. Subtelny, S. W. Eichhorn, G. R. Chen, H. Sive, D. P. Bartel, Poly(A)-tail profiling reveals an embryonic switch in translational control. Nature 508, 66–71 (2014) |
| Human | HEK293 | H3K27ac-ChIP-Seq | www.encodeproject.org | ENCSR000FCH | The ENCODE Project Consortium, The ENCODE (ENCyclopedia Of DNA Elements) Project. Science 306, 636–640 (2004) |
| Human | HEK293 | Ribo-Seq | www.ncbi.nlm.nih.gov/geo/ | GSM3566397, GSM3566398, GSM3566399 | T. F. Martinez, et al., Accurate annotation of human protein-coding small open reading frames. Nature Chemical Biology 16, 458–468 (2020) |
| Human | HEK293 | Ribo-Seq | www.ncbi.nlm.nih.gov/geo/ | GSM1887643 | A. O. Subtelny, S. W. Eichhorn, G. R. Chen, H. Sive, D. P. Bartel, Poly(A)-tail profiling reveals an embryonic switch in translational control. Nature 508, 66–71 (2014) |
| Human | HEK293 | DM-tRNA-Seq | www.ncbi.nlm.nih.gov/geo/ | GSM1624820, GSM1624821 | G. Zheng, et al., Efficient and Quantitative High-Throughput tRNA Sequencing. Nature Methods 12 (9): 835–37 (2015) |
| Human | HEK293 | hydro-tRNA-Seq | https://doi.org/10.1016/j.celrep.2017.07.029 | Table S5 | T. Gogakos, et al., Characterizing Expression and Processing of Precursor and Mature Human tRNAs by Hydro-tRNAseq and PAR-CLIP. Cell Reports 20 (6): 1463–75 (2017) |
| Fly | S2 | cytosolic polyA-RNA-Seq | www.ncbi.nlm.nih.gov/geo/ | GSM1477474 | J. L. Aspden, et al., Extensive translation of small Open Reading Frames revealed by Poly-Ribo-Seq. Elife 3, e03528 (2014) |
| Fly | ML-Dmd17-c3 | cytosolic polyA-RNA-Seq | www.encodeproject.org | ENCSR622ROA | L. P. B. Bouvrette, et al., CeFra-seq reveals broad asymmetric mRNA and noncoding RNA distribution profiles in Drosophila and human cells. RNA 24, 98–113 (2018) |
| Fly | Early embryo | cytosolic polyA-RNA-Seq | www.ebi.ac.uk/arrayexpress/ | E-MTAB-4571 | H. Li, et al., Ultra-deep sequencing of ribosome-associated poly-adenylated RNA in early Drosophila embryos reveals hundreds of conserved translated sORFs. DNA Research 23, 571–580 (2016) |
| Fission yeast | Rich growth medium (YES) | cytosolic polyA-RNA-Seq | www.ncbi.nlm.nih.gov/geo/ | GSM2805397, GSM2805401, GSM2805405 | L. Herzel, K. Straube, K. M. Neugebauer, Long-read sequencing of nascent RNA reveals coupling among RNA processing events. Genome Res. 28, 1008–1019 (2018) |
| Fission yeast |  | cytosolic rRNA-depleted RNA-Seq | www.ncbi.nlm.nih.gov/geo/ | GSM1276537 | A. O. Subtelny, S. W. Eichhorn, G. R. Chen, H. Sive, D. P. Bartel, Poly(A)-tail profiling reveals an embryonic switch in translational control. Nature 508, 66–71 (2014) |

**Table S4:** Spearman correlation coefficients with p values for Figures 2B and 3A.

**Data for Figure 2B: Correlation between codon frequencies in different ORF types**
|  | Spearman correlation coefficient |  |  |  |  |  | P value (log10) |  |  |  |  |  |
| --- | --- | --- | --- | --- | --- | --- | --- | --- | --- | --- | --- | --- |
|  | Human | Mouse | Fish | Fly | Worm | Fission yeast | Human | Mouse | Fish | Fly | Worm | Fission yeast |
| lincRNA - mRNA | 0.70 | 0.67 | 0.86 | 0.48 | 0.75 | 0.80 | -9.79 | -8.71 | -18.99 | -4.18 | -12.22 | -14.57 |
| lincRNA - control | 0.96 | 0.95 | 0.79 | 0.87 | 0.99 | 0.93 | -36.55 | -31.18 | -14.29 | -20.22 | -50.69 | -27.52 |
| lincRNA - 3UTR | 0.92 | 0.91 | 0.72 | 0.77 | 0.89 | 0.97 | -26.32 | -24.69 | -10.48 | -12.87 | -22.36 | -40.48 |
| mRNA - control | 0.62 | 0.58 | 0.62 | 0.44 | 0.80 | 0.95 | -7.45 | -6.36 | -7.32 | -3.50 | -14.60 | -32.62 |
| mRNA - 3UTR | 0.63 | 0.59 | 0.49 | 0.56 | 0.62 | 0.77 | -7.64 | -6.64 | -4.37 | -5.87 | -7.44 | -13.18 |
| control - 3UTR | 0.97 | 0.97 | 0.91 | 0.79 | 0.87 | 0.91 | -38.02 | -41.20 | -24.59 | -14.33 | -20.22 | -23.96 |

**Data for Figure 3A: Correlation between codon frequencies and tRNA abundances for different ORF types**
|  | Spearman correlation coefficient |  |  |  |  |  | P value (log10) |  |  |  |  |  |
| --- | --- | --- | --- | --- | --- | --- | --- | --- | --- | --- | --- | --- |
|  | Human | Mouse | Fish | Fly | Worm | Fission yeast | Human | Mouse | Fish | Fly | Worm | Fission yeast |
| mRNA | 0.58 | 0.53 | 0.46 | 0.58 | 0.58 | 0.46 | -5.83 | -4.79 | -3.64 | -5.88 | -5.98 | -3.73 |
| top-cyto-mRNA | 0.58 |  |  | 0.59 |  | 0.59 | -5.82 |  |  | -6.08 |  | -6.13 |
| lincRNA | 0.29 | 0.26 | 0.38 | 0.13 | 0.14 | 0.10 | -1.58 | -1.31 | -2.53 | -0.50 | -0.54 | -0.37 |
| cyto-lincRNA | 0.31 |  |  | 0.12 |  | 0.10 | -1.77 |  |  | -0.42 |  | -0.34 |
| intronic&intergenic | 0.27 | 0.18 | 0.30 | 0.13 | 0.17 | 0.32 | -1.39 | -0.77 | -1.69 | -0.48 | -0.71 | -2.07 |
| 3'UTR | 0.32 | 0.18 | 0.21 | 0.16 | 0.07 | 0.09 | -1.88 | -0.78 | -0.97 | -0.64 | -0.21 | -0.29 |
| lincRNA (first ORF) | 0.30 | 0.27 | 0.33 | 0.15 | 0.13 | 0.06 | -1.73 | -1.44 | -1.99 | -0.61 | -0.51 | -0.20 |
| cyto-lincRNA (first ORF) | 0.33 |  |  | 0.10 |  | 0.10 | -2.04 |  |  | -0.36 |  | -0.34 |
| intronic&intergenic | 0.25 | 0.20 | 0.28 | 0.06 | 0.16 | 0.23 | -1.24 | -0.89 | -1.38 | -0.19 | -0.68 | -1.13 |
| 5'UTR (first ORF) | 0.23 | 0.29 | 0.25 | 0.12 | 0.32 | 0.09 | -1.09 | -1.64 | -1.28 | -0.46 | -1.89 | -0.29 |

